# Molecular heterogeneity of *C. elegans* glia across sexes

**DOI:** 10.1101/2023.03.21.533668

**Authors:** Maria D. Purice, Elgene J.A. Quitevis, R. Sean Manning, Liza J. Severs, Nina-Tuyen Tran, Violet Sorrentino, Manu Setty, Aakanksha Singhvi

## Abstract

A comprehensive description of nervous system function, and sex dimorphism within, is incomplete without clear assessment of the diversity of its component cell types, neurons and glia. *C. elegans* has an invariant nervous system with the first mapped connectome of a multi- cellular organism and single-cell atlas of component neurons. Here we present single nuclear RNA-seq evaluation of glia across the entire adult *C. elegans* nervous system, including both sexes. Machine learning models enabled us to identify both sex-shared and sex-specific glia and glial subclasses. We have identified and validated molecular markers *in silico* and *in vivo* for these molecular subcategories. Comparative analytics also reveals previously unappreciated molecular heterogeneity in anatomically identical glia between and within sexes, indicating consequent functional heterogeneity. Furthermore, our datasets reveal that while adult *C. elegans* glia express neuropeptide genes, they lack the canonical *unc-31/CAPS*-dependent dense core vesicle release machinery. Thus, glia employ alternate neuromodulator processing mechanisms. Overall, this molecular atlas, available at www.wormglia.org, reveals rich insights into heterogeneity and sex dimorphism in glia across the entire nervous system of an adult animal.

## INTRODUCTION

The nervous system is a complex network of neurons and glia, where glial cells play critical roles in modulating neuronal properties and animal behavior. However, due to the lack of tools to study different glial subtypes, our understanding of the cellular and molecular level interactions between glia and neurons remains incomplete.

The nematode *Caenorhabditis elegans* is a powerful model organism for studying glia-neuron interactions (Shaham, 2010; Singhvi and Shaham, 2019). Descriptions of the connectomes for both adult sexes and a late larval transcriptional atlas of hermaphrodite neurons have allowed for the study of circuits underlying a wide range of behaviors. However, our understanding of the glial cells that interact with these neurons and circuits at the cellular and molecular level remains incomplete, in part due to the lack of tools to study different glial subtypes in this organism.

High throughput technologies like single cell and single nuclei RNA-sequencing (scRNA-seq and snRNA-seq) have ushered new avenues for exploring the diversity of cells types in the brain. External reference databases such as the Human Cell Atlas, Mouse Cell Atlas, and other cell type specific databases such as brainRNAseq.org (mammalian glia) and cengen.org (*C.elegans* neurons) have facilitated the linkage of gene expression to cell class and function. These studies also uncover insights into the complexity and dynamics of cell states longitudinally (Lago-Baldaia et al., 2022; Setty et al., 2019; Soreq et al., 2017). However, across species, identification of molecular classifications of glial cells has been more challenging compared to neurons, due to lack of cell type-specific and universal markers (Herculano-Houzel and Dos Santos, 2018; Zhang et al., 2014; Zhang and Barres, 2010).

The nervous system of the adult *C. elegans* is composed of 302 neurons and 56 glia in the hermaphrodite, of which 50 glia are neuroectoderm-lineage derived and are found in sense- organs (Sulston and Horvitz, 1977; Ward et al., 1975). Within each sense organ, dendrites of one or more bipolar sensory neurons traverse a channel created by a single **sh**eath (sh) glia and one or more **so**cket (so) glia (Bacaj et al., 2008; Ward et al., 1975). Sheath glia are anatomically defined as having anterior membranes that surround the receptive ending at the dendrite tip of sensory neurons, while socket glia interface between epithelia and sheath glia, allowing the sensory dendrites to extend into and sample the environment. One class of sheath glia, CEPsh, also has posterior processes that interact with the brain neuropil of the animal, where its functions are considered analogous to vertebrate astrocytes (Katz et al., 2018; Singhvi and Shaham, 2019). Six glia in the animal, the GLR, are mesodermal-derived (Altun and Hall, 2003). They extend sheath-like projections into the nerve ring and make gap junctions with at least one neuron to establish its axon specification (Stout Jr. et al., 2014).

Anatomically, the hermaphrodite animal has 24 sheath and 26 socket glia across 7 sense organs (Figure 1A, B). The *C. elegans* male nervous system has 389 neurons and 92 glia, of which 86 glia are neuroectoderm-lineage derived (Fig 1A, B). These 86 glia further subclassify into 10 sheath/socket subtypes, as well as the male-specific ray structural glia-like cells, which are proposed to have properties of both sheath and sockets (Lints and Hall, 2005). However, the majority of these glial cell types remain understudied at molecular detail, and for some, molecular markers are not available. It is also unclear if the anatomical sheath or socket glia classes implies functional equivalence. Finally, the extent of heterogeneity across glia in this animal has not been explored.

**Figure 1.**
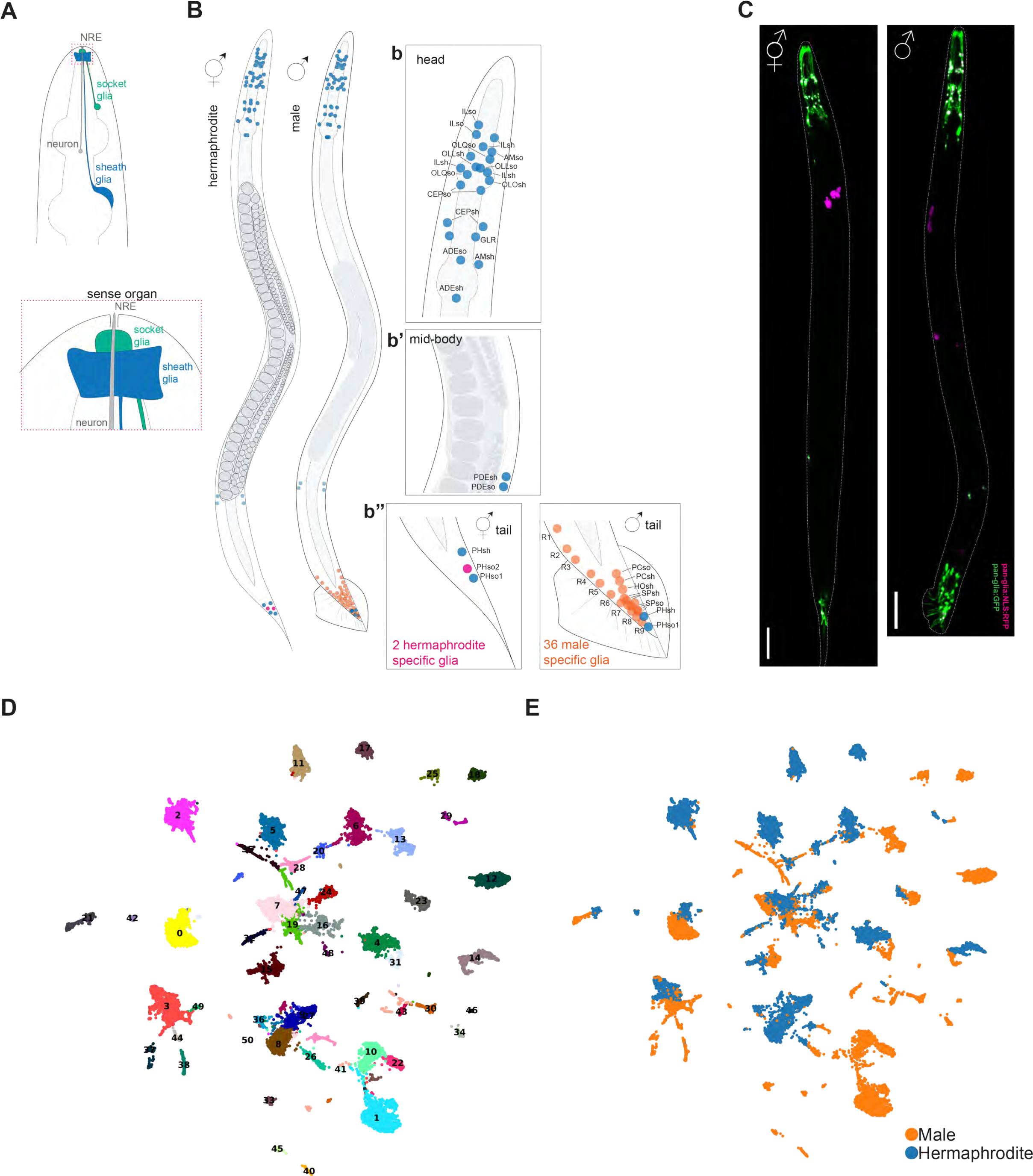
Anatomical and molecular characterization of adult C.elegans glia in hermaphrodites and males using pan-glial transcriptional reporter *miR-228*. (**A**) Schematic example of a *C.elegans* sense organ or sensilla, consisting of one or more sensory neurons (gray) that associate with one socket (green) and one sheath (blue) glia. Below is a close-up of the nose tip (dotted magenta box) showing interactions between the neuron and two glia. (**B**) Schematic representation of adult hermaphrodite and male showing sex shared (blue), hermaphrodite-specific (magenta), and male-specific (orange) glial nuclei. Close-up of the head (b), mid-body (b’) and hermaphrodite and male tails (b”) and glia within the region. (**C**) Z-stack projection and stitched tiles depict adult hermaphrodite (left) and male (right) expressing pan-glial cytoplasmic GFP and nuclear localized RFP. Animals are also expressing co-injection marker coelomocyte RFP. Scale bars = 50μM. (**D**) UMAP of 51 non-batch corrected clusters from day one adult hermaphrodites and males. (**E**) UMAP of 51 non-batch corrected clusters from day one adult hermaphrodites and males showing sex contribution to each cluster (male samples = orange; hermaphrodite samples = blue). Genotypes: Figure 2C: *him-5; PmiR-228:GFP; PmiR-228:nls:RFP, Punc-122:RFP*.

Evidence in mammals suggest that glial sexual dimorphism contributes to the development and function of neurons (Nguon et al., 2005; Schwarz and Bilbo, 2012; Simerly, 2002). Similar to vertebrates, *C.elegans* nervous system has some anatomically dimorphic glia between sexes.

Specifically, adult males have an additional 89 neurons and 36 glia. Of the 92 total glia between hermaphrodites and males, 54 are shared by both sexes, two are hermaphrodite specific and 36 are male specific. The adult hermaphrodite has two glia (**ph**asmid **so**cket 1 (PHso1)) that transdifferentiate into neurons in the male and are therefore absent (Molina-García et al., 2020). All male specific glia reside in four sensilla groups in the male tail, many of which control male mating-related behaviors (Sulston et al., 1980). Further, some of the sex-shared glia interact with sexually dimorphic neurons and circuits in the brain neuropil of the animal. For example, the astrocyte-like **ce**phalic **sh**eath (CEPsh) glia associates with the sensory endings of dopaminergic CEP neurons in both sexes, but additionally also with the male-specific cholinergic CEM neuron in the male (Sammut et al., 2015). Similarly, the sex-shared **am**phid **so**cket (AMso) glia divides to generate the MCM neuron only in the male (Sammut et al., 2015). Lastly, PHso1 glia transdifferentiates only in the male (Molina-García et al., 2020). Whether these sex-dimorphisms imply different functional profiles for any of these glia is unknown.

To define glial heterogeneity and sex dimorphism of an adult animal nervous system, we performed snRNA-seq on adult hermaphrodite and male glia. Our analyses identified 32 distinct gene expression profiles, of which 1 was specific to hermaphrodites, 9 were specific to males, and the remaining 22 profiles consisted of cells from both sexes. Using iterative computational methods, we identified novel glial subtype-specific markers and have validated these *in vivo* using transcriptional reporters. Furthermore, using machine learning models, we identified sheath-specific, socket-specific, and pan-glial markers. Our findings revealed previously unknown heterogeneity within sex-shared glia, with some sex-shared glia exhibiting unanticipated molecular differences between sexes, such as PHsh. We also see instances of molecular convergence for glia that does not track predictions based on anatomy, for example male SPsh/so, Hosh/so, and PCsh/so. Lastly, although several neuropeptides were expressed in glial cells, the genes for canonical dense-core vesicle machinery release were conspicuously downregulated. As glia do express neuropeptides (Frakes et al., 2020), this implies that they employ an alternative neuromodulator packaging mechanism to do so. Thus, this molecular atlas and functional validation studies reveal previously unappreciated insights into the biology and heterogeneity of glia in the simple nervous system of *C. elegans*. This also creates tools to investigate the organizational principles underlying glial functions in this animal model.

## RESULTS

### Anatomical and molecular characterization of adult *C. elegans* glia across sexes

To identify and isolate *C. elegans* glia, we anatomically characterized the only reported pan-glial promoter, that of the microRNA *miR-228* (Fung et al., 2020; Wallace et al., 2016). We examined a transcriptional fluorescent reporter in both sexes in day one adult animals (Day 1 Ad) and observed expression in glia throughout the animal body-plan: head, midbody and tail (Figure 1C). We also observed expressions in the excretory canal, vulva, rectum and hypodermis in the tail (Figure S1A). We did not observe expression in early embryos prior to egg-laying, in the germ line, or in the mesoderm-derived GLR glia (Figure S1B, C).

To identify the transcriptional profile of adult hermaphrodite and male glia, we performed single nuclear RNA-sequencing (snRNA-seq) on Day 1 Ad hermaphrodites and males. Prior *C. elegans* single cell RNA-seq (scRNA-seq) data has shown that even after FACS, there can be persistent non-specific cell contamination (Taylor et al., 2021); and anatomically glia and neurons have intertwined processes but physically distant cell bodies. We therefore chose snRNA-seq, with the aim that restricting analyses to glial nuclei will likely avoid contamination by physically associated neuronal cell material. We labeled glial cell nuclei by expressing nuclear RFP under P*miR-228.* After enzymatic dissociation of the animal tissue into single nuclei suspension [refer Methods for adapted protocol based on (Kaletsky et al., 2016)], we used **f**luorescence-**a**ctivated **c**ell **s**orting (FACS) to select RFP+ nuclei. As we confirmed that P*miR-228* does not express in the early embryo or germ cells, this transgene allowed us to avoid contamination with germ cells or early embryonic cells.

Obtaining sufficient material for snRNA-seq on male glia is challenging because wild type *C. elegans* reproduces primarily as self-fertilizing hermaphrodites with males occurring at population frequencies of only 0.1% (Chasnov and Chow, 2002). To obtain a large number of cells from adult males, we used a combination of three methods. We first introduced a *him-5* mutation, which induces aneuploidy and generates ∼30% XO males in the population (Cook et al., 2019). Secondly, we labeled our transgene with mVenus driven under the *mig-24* promoter. P*mig-24* expresses in two cells in the hermaphrodite lifelong distal tip cells. In the male, this promoter drives expression only in the analogous linker cell, which undergoes cell death at the L4-to-adult transition (Malin et al., 2016). Thus, enriched *him-5* adult males can be identified by loss of P*mig-24*:mVenus fluorescence in the animal. Lastly, we took advantage of the fact that males and hermaphrodite animals are of different sizes. Thus, we first washed Day 1 Ad *him-5* animals using sieves to separate out adult hermaphrodites. We then used a COPAS worm sorter to select animals based on size and lack of mVenus reporter expression. Once obtained, these animals were subjected to identical cell dissociation and FACS protocols as hermaphrodites.

For both sexes, we validated enrichment of nuclei after FACS in two ways. One, we stained and imaged RFP+ and RFP- sorted nuclei with the DNA-binding dye propidium iodide (PPI) (Crowley et al., 2016). Although both RFP+ and RFP- fractions were stained with PPI, only the RFP+ fraction contained RFP signal (Figure S1D), confirming enrichment of glial nuclei in the sample. Two, we performed quantitative real time PCR on FACS sorted RFP+ nuclei and observed enrichment of known glial genes *F53F4.13, kcc-3, and ptr-10* (Fung et al., 2020) and a depletion of neuronal gene *unc-119* (Figure S1E), further confirming enrichment of glial transcripts.

After sequencing using the 10x Genomics platform, we removed empty droplets using EmptyDrops methods implemented in the R package DropletUtils. Through iterative verifications, we chose 200 UMIs as the optimal cutoff to define empty droplets. The final dataset contained 31410 total cells, 16887 hermaphrodites and 14723 males with a median of 939 UMIs and 440 genes/cell. We applied Leiden clustering and used the **U**niform **M**anifold **A**pproximation and **P**rojection (UMAP) dimensional reduction algorithm to uncover 51 clusters of which 10 were hermaphrodite only, 22 male only, and 19 were both (Figure 1D, E).

### Iterative computational and experimental validation to identify glial clusters

As the gene expression profile of adult *C. elegans* glia is not known, we first used existing glial markers from literature (Cao et al., 2017; Fung et al., 2020; Packer et al., 2019; Wallace et al., 2016) to identify clusters for known glia. We were confidently able to identify one cluster consisting of the AMsh and PHsh glia, two related glial cell types (Figure 2A, Figure S2A) by high expression of known genes including*, F53F4.13*, *T02B11.3, fig-1, and vap-1* (Fung et al., 2020; Wallace et al., 2016). Thus, the molecular profile of at least these glial cells are reliably identified in our dataset. Interestingly, the AMsh/PHsh cluster is made of two subclusters with only the lower one expressing *vap-1*, a known AMsh-specific gene that is absent in PHsh (Figure 2A). This implies that although the AMsh/PHsh are transcriptionally similar and form one cluster, the cells in the upper cluster are PHsh and the *vap-1*+ cells are AMsh. To further validate that cluster 14 is indeed the AMsh/PHsh glia, we made transcriptional reporters with upstream regulatory regions for two highly enriched genes, *ZK822.4* and *far-8* that had not yet been identified as AMsh/PHsh glia-enriched. Indeed, the transcriptional reporter for *ZK822.4* showed specific expression in the AMsh/PHsh glia in both hermaphrodites and males, however the *far-8* reporter showed mostly hermaphrodite tail expression (Figure 2B, C). While picking P*far-8*: GFP animals for imaging, we observed that some adults only lacked AMsh or PHsh expression, but many of the young larvae had high expression of GFP in both AMsh and PHsh. We thus picked recently hatched L1 animals that had expression of AMsh/PHsh in the tail and scored their expression in Day 1 Ad. We observed that most hermaphrodites retain expression only in the PHsh while males lose expression in both cells (Figure 2C, D). These results suggest that *far-8* is temporally and differentially expressed.

**Figure 2.**
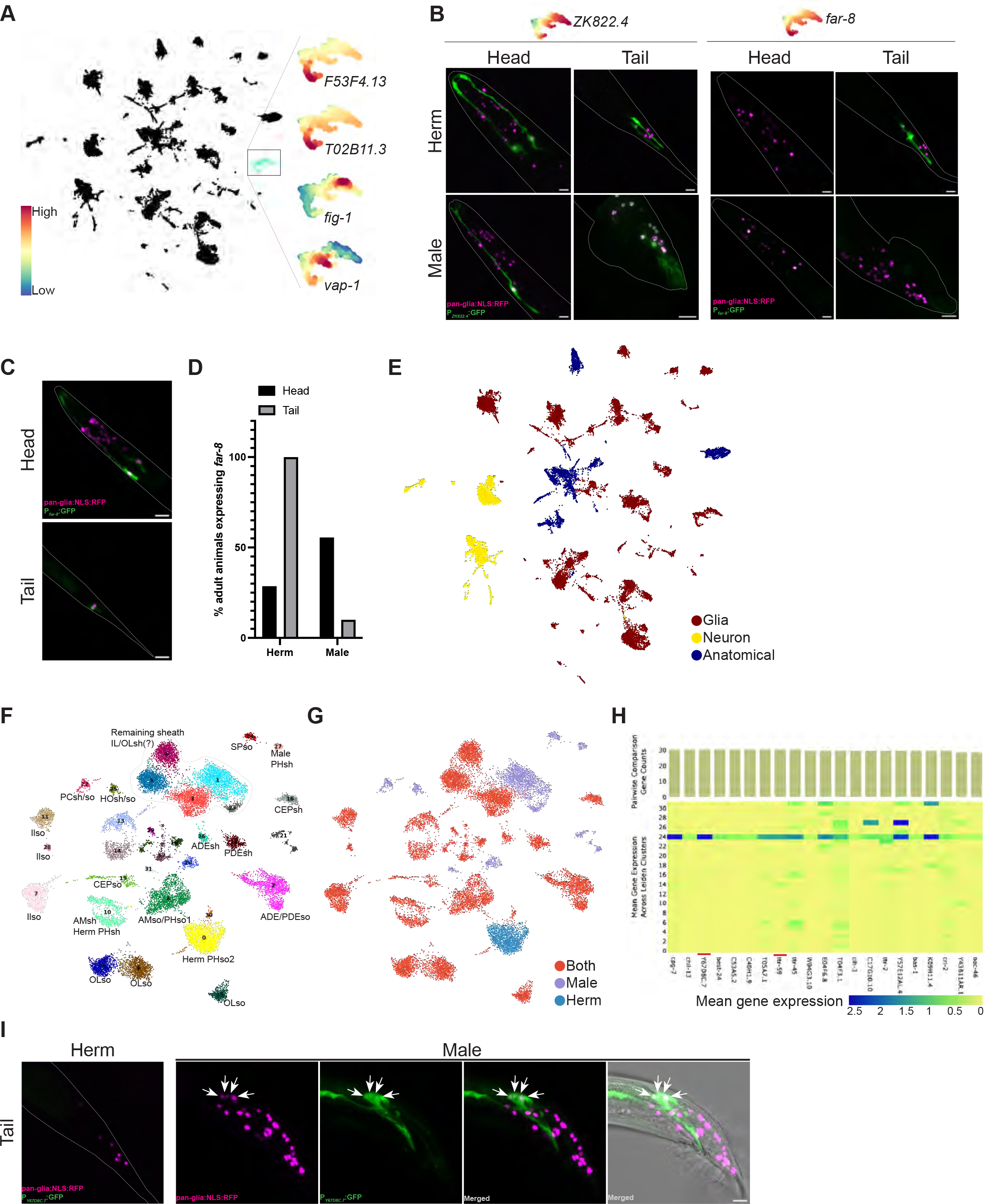
Validation of cluster identities. (**A**) UMAP projection depicting expression of known AMsh/PHsh genes (*F53F4.13, T02B11.3, fig-1, vap-1)* within Cluster 14. Heatmap shows expression level. (**B**) Z-stack projections of adult hermaphrodite and male heads and tails with dotted outlines. Mean gene expression of each gene within cluster 14 shown at the top. Left side panels shows a transcriptional reporter for *ZK822.4* depicting expression in the AMsh/PHsh glia in both sexes. Right side shows the *far-8* reporter displaying mostly hermaphrodite tail expression. (**C**) Z-stack projections of L1 animal showing expression of transcriptional reporter for *ZK822.4* in AMsh/PHsh. (**D**) Graph depicting the percentage of adults that retained expression in AMsh/PHsh in each sex. Animals like the representative in (**C**) were picked as L1 larvae if they had expression in both the head and tail (i.e. AMsh and PHsh) and then were scored again as day one adults. N=18 males and 21 hermaphrodites. (**E**) Non-batch corrected UMAP depicting 34 glial cell clusters (dark red), 8 neuronal clusters (yellow), and 8 anatomical clusters (dark blue). (**F**) UMAP of 32 batch corrected glial only clusters and their identifies. (**G**) UMAP of 32 batch corrected glial only clusters depicting sex specificity: 1 hermaphrodite- specific cluster (blue), 9 male-specific clusters (lavender), and 22 sex-shared clusters (red). (**H**) Pairwise comparison analysis for all male cluster 24 showing expression of genes (x-axis) per cluster (y-axis). Counts per gene for each sample shown at the top. The higher the number to total number of clusters (i.e. 32), the more unique the expression of the gene. Genes *Y67D8C.7* and *ttr-59* (red dashes) were chosen due to their high and unique expression. (**I**) Z-stack projections of the *Y67D8C.7* transcriptional reporter. Hermaphrodite tail shown as merged. Male tail shown as individual and merged channels. Arrows point to the four *miR-228* RFP*+* nuclei that co-localize with the GFP+ cells projecting into the spicule. Genotypes: Figure 2B: *him-5; PZK822.4:GFP, Punc-122:GFP; PmiR-228:nls:RFP, Punc-122:RFP* & *him-5; Pfar-8:GFP, Punc-122:GFP; PmiR-228:nls:RFP, Punc-122:RFP*. Figure 2C, D: *him-5; Pfar-8:GFP, Punc-122:GFP; PmiR-228:nls:RFP, Punc-122:RFP*. Figure 2I: *him-5; PY67D8C.7:GFP, Punc-122:GFP; PmiR-228:nls:RFP, Punc-122:RFP*. All scale bars = 10μM.

Surprisingly, many known genes were not highly expressed in the adult dataset, including *ptr-10,* known to express broadly across many glia. Although transcriptional reporters for *ptr-10* show glial expression in adult animals, the expression levels of *ptr-10* transcript in adults are barely detected, even in the sorted glial fraction by snRNAseq or qPCR (Figure S1F, Figure S2C). This suggests that developmentally expressed genes and extra-chromosomal array reporters may not be reliable in predicting endogenous transcriptional gene expression in adult glia.

The *miR-228* reporter labels other non-glial cells. To remove these we used the available scRNA-seq dataset from *C. elegans* larval stage L4 (Taylor et al., 2021). We separated 34 glial cells clusters (21082 cells, 67.12%) from non-glial cells that were separated into two categories, 8 neuronal clusters (5055 cells, 16.09%), and 9 anatomical clusters (5273 cells, 16.79%) (Figure 2E). Interestingly, some of the neuronal specific clusters were composed of male cells and their characterization awaits further studies (Figure S2D). We performed hierarchical clustering on the three subclusters of cells (glia, neurons, anatomical) and found that sheath glia are independently grouped while the socket glia are similar to the neuronal and anatomical clusters (Figure S2E).

On the subset data, we then performed batch correction and reiterated dimensionality reduction to generate a new UMAP of batch corrected glia only clusters (Figure 2F). All subsequent analyses described are on these data. Unsupervised clustering of these revealed 32 glial clusters and each cluster was then assigned a label denoting sex-specificity (Figure 2G). If a cluster was greater than 95% for a certain sex, it was deemed specific to be either hermaphrodite or male, otherwise it was labeled both (Figure 2G, S2F). This analysis uncovered 1 hermaphrodite-specific cluster, 9 male-specific clusters, and 22 sex-shared clusters (Figure 2G). We further used a machine learning model trained on either male or hermaphrodite batch corrected data to predict sex specificity. When the model was trained on male specific data, we observe a 1 to 1 mapping to our manually assigned sex labels. However, when the data was trained on hermaphrodite specific data, we observed the expected a 1 to 1 mapping with a case with a few exceptions (clusters 1, 3, 30) (Figure S2G). Thus, for reasons not yet clear, the male glia dataset is a more robust training set for machine learning algorithms.

We also considered that some of the variance observed in our original datasets might reflect biological, rather than technical, variability. Therefore, we also independently performed analogous verification on non-batch corrected data (Figure 1D, E). However, this led to unreliable downstream analytics (see Methods section), with some original clusters being merged (Figure 2F, S2H). We also observed partial non-equivalence in pairwise comparison analyses in some clusters between batch-corrected and non-corrected datasets. Therefore, while it remains possible that the molecular variance in non-batch corrected datasets between sexes is biologically relevant, we focused on batch-corrected datasets as the first pass conservative approach.

Next, we sought to reveal the identity of the remaining clusters and performed pairwise comparison analysis to uncover **u**niquely **e**xpressed **g**enes (UEG) within each cluster. To functionally validate these predictions, we generated transgenic animal strains where promoter of a UEG drove GFP as a transcriptional reporter in a background of pan-glia labeling using the P*miR-228*:NLS:RFP strain. This approach enabled us to assign cell-specific anatomical identity to each glial cluster, as well as identify multiple novel markers for glia.

For example, in the pairwise comparison for male specific cluster 24, we selected two UEGs: *Y67D8C.7* and *ttr-59* for *in vivo* validations (Figure 2H). A transgenic reporter strain of P*Y67D8C.7*:GFP did not show glial expression in the hermaphrodite heads or tails or the male heads (Figure 2I). However, GFP expression was consistently observed in four syncytial cells that colocalize with RFP and project into the spicule (Figure 2I). This led us identify that *Y67D8C.7* expresses in the **sp**icule **so**cket (SPso) glia. Similar to P*Y67D8C.7*:GFP, P*ttr-59*:GFP also expressed in the four syncytial cells that colocalize with RFP and project into the spicule (Figure S2I). These data identify the male specific cluster 24 as that for SPso glia.

Analogous methodological pipeline allowed us to similarly assign cell-specific identities to 24/32 clusters on the glial UMAP (Figure 2F). All cell expression and gene ID used to validate each glial cluster, along with the upstream regulatory region used to drive the transcriptional reporter, is tabulated in entirety as (Table S1).

For clusters 1, 3, 5, 6 and 23 we were unable to find UEGs or transcriptional reporters that expressed in RFP+ glia, suggesting high overlap in their gene-expression profiles. By process of elimination we identify these as **i**nter **l**abial and **o**uter **l**abial **sh**eath (IL/OLsh) glia (Figure 2F, outline). Despite their high level of similarity in gene expression, the presence of multiple clusters indicates that these glia are likely non-identical.

During the *in vivo* validations, we observed ectopic expression of GFP in other cell types, in addition to the glia, suggesting that while the UEG may differentiate with other glia, it does not indicate exclusive expression. For example, we anticipated cluster 17 to be **cep**halic **sh**eath (CEPsh) glia based on expression of *hlh-17*, a previously identified as a CEPsh-expressing gene [Fung]. Validation of cluster 17 with a novel GFP transcriptional reporter (P*Y71H10B.1*:GFP) showed expression in CEPsh as expected and also in a second group of glia (Figure S2J). Apart from identifying cluster identities, these overlaps can also provide valuable insights for future research on the potential functional similarities suggested by these expression patterns.

### Hierarchical unsupervised clustering identifies heterogeneity in glial subpopulation signature profiles

Each sense organ contains both sheath and socket glia (Figure 1A). Given that most glia cluster individually, we next asked whether glia of a single organ cluster together, or whether they cluster functionally as sheath or socket glia. For this, we performed cosine similarity using highly variable genes and found that the glial clusters segregate into two groups (Figure 3A).

**Figure 3.**
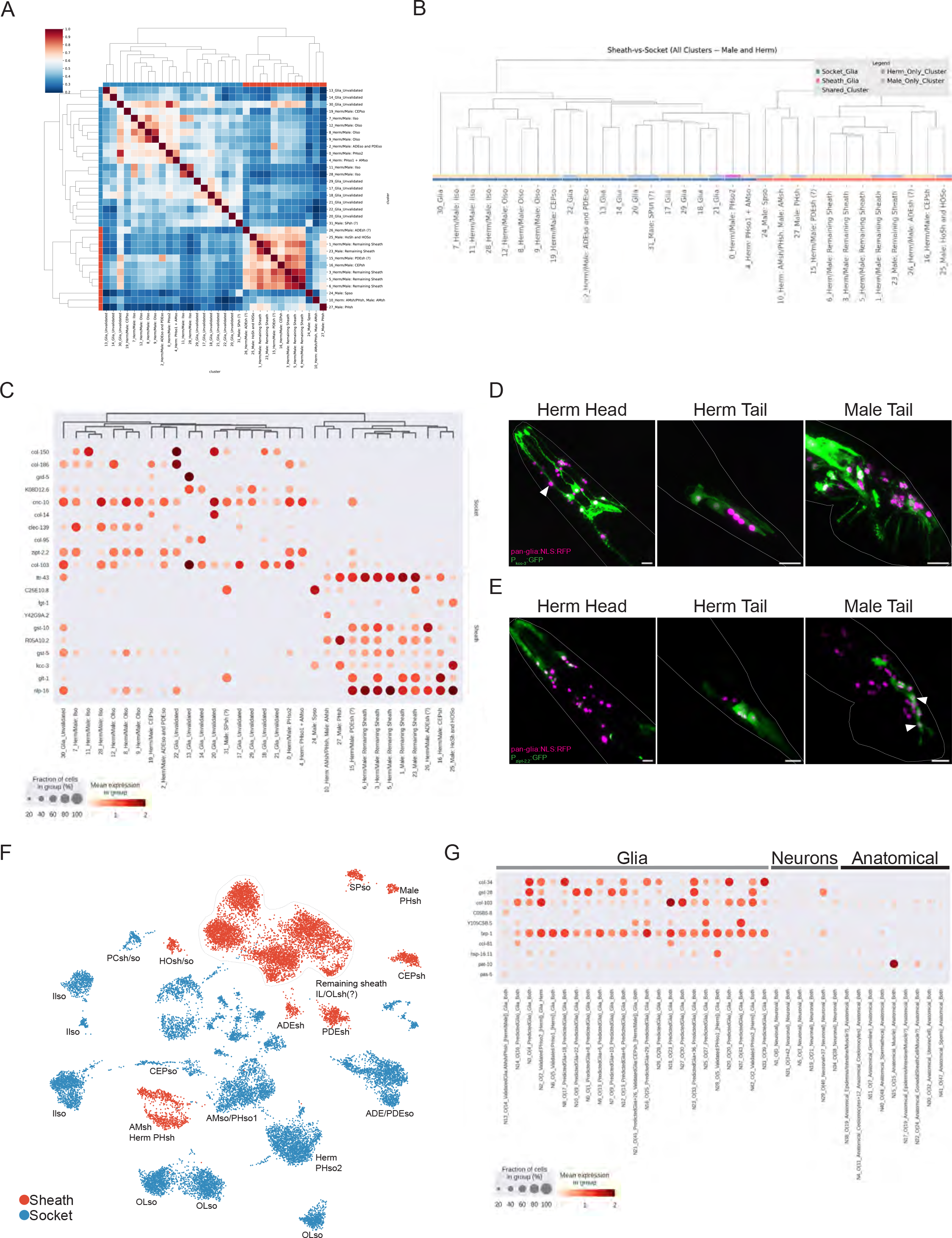
Unsupervised clustering identifies markers for populations of glia. (**A**) Cosine similarity using highly variable genes shows relationship between clusters. (**B**) Hierarchical clustering shows separation of clusters into two groups on a dendrogram. (**C**) Binary classifier machine learning model’s top 10 candidate genes for sheath and socket markers. (**D**) Z-stack projection of the *kcc-3* transcriptional reporter in hermaphrodite head/tail and male tail shows expression in sheath glia. Arrowhead depicts GFP- AMso glial nuclei. (**E**) Z-stack projection of the *zipt-2.2* transcriptional reporter in hermaphrodite head/tail and male tail shows expression in socket glia. Arrowheads depicts GFP+ in PHso2 and PHso1-derived PHD neuron. (**F**) UMAP of batch-corrected glial only clusters sheath/socket identity based on molecular identity. (**G**) Binary classifier machine learning model’s top 10 candidate genes for pan-glia markers. Genotypes: Figure 3D: *him-5; Pkcc-3:GFP; PmiR-228:nls:RFP, Punc-122:RFP*. Figure 3E: *him-5; Pzipt-2.2:GFP, Punc-122:GFP; PmiR-228:nls:RFP, Punc-122:RFP*. All scale bars = 10μM.

Additionally, hierarchical clustering revealed again the separation of clusters into two groups on a molecular dendrogram (Figure 3B). The cluster identity we had established led us to recognize these groups as the sheath and socket classes (Figure 3B). This provides independent molecular validation that sheath glial functions across sense organs are molecularly more related to each other than to the socket glia within their own sense-organs that interact with the same neuron.

This molecular convergence also does not track development because sheath glia, socket glia and neurons develop in intermingled lineages. Thus, sheath and socket glia likely have subclass identity selector transcriptional programs, similar to how different neuron subclasses are specified (i.e. cholinergic, glutamatergic) (Hobert, 2010).

To probe this further, we used a data driven approach to uncover novel markers for the two classes of glia. We used the batch corrected data that contain either male- or hermaphrodite-specific clusters separately and used the sheath/socket label assignment to distinguish each cluster in the dataset. To identify candidate markers for sheath and socket cells, **m**achine **l**earning **m**odel (MLM) was trained using imputed gene expression values. The MLM is a binary classifier, meaning that it was designed to distinguish between sheath and socket cells based on their gene expression patterns. By running the MLM on the gene expression data, the resulting markers that were identified are likely to be associated with either sheath or socket cells.

The resulting candidate markers for sheath and socket was identified by training a binary classifier MLM to distinguish sheath and socket cells given imputed gene expression values. After the MLM was trained on the male dataset, the top 10 unimputed features or genes that were most strongly associated with either sheath or socket cells were then displayed (Figure 3C, Figure S3A, S3B). This analysis shows that sheath versus socket distinction is maintained globally, with distinct sheath and socket signatures that generalize across both sexes. The strongest and most inclusive markers for sheath glia were *kcc-3* and *ttr-43*, while *zipt-2.2 and cnc-10* were the most representative genes for socket expression (Figure 3C).

It has been previously shown that *kcc-3* is expressed in multiple glia including AMsh and possibly a subset of socket glia (Fung et al., 2020; Singhvi et al., 2016; Yoshida et al., 2016).

Since our binary classifier model found *kcc-3* as a sheath glia specific gene, we characterized *kcc-3* expressing cell types using a GFP transcriptional reporter. When crossed to the pan-glial nuclear RFP line, we observed expression in AMsh glia as expected, as well as CEPsh, and PHsh glia in the hermaphrodites, and in cells around the first pharyngeal bulb (Figure 3D). Similar expression was seen in adult male heads, as well as expression in ray structural glia and additional structures expressed in the tail (Figure 3D). We do not observe *kcc-3* expression in the **ph**asmid **so**cket (PHso) or **am**phid **so**cket (AMso) glia (Figure 3D, arrowhead). Taken together, these results confirm MLM modeling to reveal *kcc-3* as a pan-sheath marker.

Next, we performed similar validation studies on the MLM-predicted socket marker *zipt-2.2.* We detected expression in socket cell subtypes in the head and in the tail (Figure 3E). As expected for a socket marker, we observed expression in the sex distinct PHso glia. In the hermaphrodite tail *zipt-2.2* is expressed in the PHso1 and PHso2, while in males, we observe expression in the PHso2 and PHD neurons (Figure 3E, arrowheads). While examining the *zipt-2.2* gene expression, we found that the reporter construct showed mosaic expression, as well as expression in non-glia cells.

Our hierarchical clustering dendrogram also reveals interesting molecular phylogeny between some glia. For example, the three ILso and OLso clusters appear in their own clades, across three separate clusters, suggesting closely related molecular profiles and functions (Figured 3B, left branch). In addition, we unexpectedly uncovered that the transcriptional identities of some male-specific glia does not track prediction based on anatomical identity. For example, the SPso glia groups with the sheath glia subclass and **sp**icule **sh**eath (SPsh) glia groups with socket glia subclass (Figure 3B). Meanwhile **p**ost **c**loacal **sh**eath/**so**cket (PCsh/so) coalesce into a combined cluster which groups with the socket-glia class, while **ho**ok **sh**eath/**so**cket (HOsh/HOso) form a combined cluster that groups with the sheath glia class (Figure 3B). Based on the UMAP visualization, we observed that the clusters of glial cells classified as either sheath or socket tend to group together on the map, with one notable exception – the AMsh/PHsh cluster was found to segregate with the sockets rather than with the other sheath cells (Figure 3F).

Finally, to date no gene has been identified that is expressed in all of glial cells besides *miR-228*, which itself is not glia-specific. Therefore, using similar MLM methods, we took a data driven approach to uncover a pan-glial markers or molecular signature profiles from our datasets. We identified a core set of candidate genes on batch corrected data that are enriched in glial cells: *brp-1, col-34* and *col-103* (Figure 3G). To confirm the identity of these candidate pan-glial signature genes, we utilized an independently published atlas of all cells in *C. elegans* during various stages of adulthood (Day 1, 3, 5, 8, 11, and 15) as a test dataset (Roux et al., 2022). This aging atlas includes detailed information on the gene expression patterns of all cells in *C. elegans* during aging (Roux et al., 2022). First, we validated that the cells captured in our dataset correspond to those found in Day 1 Ad dataset and uncovered that indeed the three subtypes of cells (i.e. glia, neurons, anatomical) captured in our RNA-seq correspond to similar subtypes captured by Roux AE et al (Figure S3C). We next derived a pan-glia signature score for our top 6 pan-glial genes (Figure S3D). Encouragingly, the pan-glial signatures colocalize with preexisting glial labels in this test dataset. However, it also identified mismatches, where it called some clusters as “glia” that had been labeled in that study as “hypodermis” or “excretory”. As the glial cells labeled previously in this dataset as “glia” account only for a fraction of all glia, we speculate that some of the clusters labeled in this dataset as “hypodermis” or “excretory” may in fact be glia, as identified by our pan-glial signature profiling. Lastly, through these analyses, we note that the pan-glial signatures stayed consistent across animal age, with only subtle changes in some glial clusters across time (Figure S3D). As a result, by overlaying our MLM modeling with these datasets, we can quickly and accurately identify specific molecular changes that occur with aging in animals. Taken together, these results indicate that the molecular signatures we identified for pan-glia, and likely socket and sheath glia subclasses, are broadly able to accurately identify glia in independently obtained snRNA-seq datasets.

### Molecular validations reveal glial sex dimorphism

Anatomical sex differences in glia are well described between hermaphrodite and males, but correlation of anatomy with glial molecular identities for sex-shared or sex-specific glia has not been explored. The datasets and validations we have obtained so far enable us to directly investigate this. Our data reveal 1 hermaphrodite-specific, 9 male-specific and 22 sex-shared glia clusters (Figure 2G).

We first examined the sex-shared CEPsh glia, which interact with receptive-endings of the CEP neurons and axons of neurons projecting within the nerve ring in both sexes. In males, however, CEPsh glia also interact with male specific CEM neurons(Singhvi and Shaham, 2019; Ward et al., 1975). We found only one shared CEPsh glial cluster in our dataset, consisting of almost equal number cells from each sex (Figure S2F, cluster 18). The lack of sex specific CEPsh glia clusters suggests that interaction with the CEM does not alter the molecular profile of this glia (Figure 2F).

At the L4 larval stage, the AMso glia in males give rise to the MCM neurons, while male PHso1 trans-differentiate into the PHD neuron (Molina-García et al., 2020; Sammut et al., 2015). The PHso1 and PHso2 cells exhibit different characteristics and connect to different cells, with PHso1 wrapping around the tip of neuronal dendrites and connecting to PHsh and PHso2, and PHso2 connecting to the hypodermis and PHso1 but not to PHsh (Hall, 1977; Sulston et al., 1980). We therefore expected AMso, PHso1, and PHso2 to each have sex-specific clusters.

However, in our adult glia datasets, both AMso and PHso1 glia coalesce into one cluster, as validated by the transcriptional reporter for *Y52E8A*.*3* (Figure 4A, S4A). We infer that despite the developmental fates in larva, adult AMso and PHso1 have similar transcriptional profiles.

**Figure 4.**
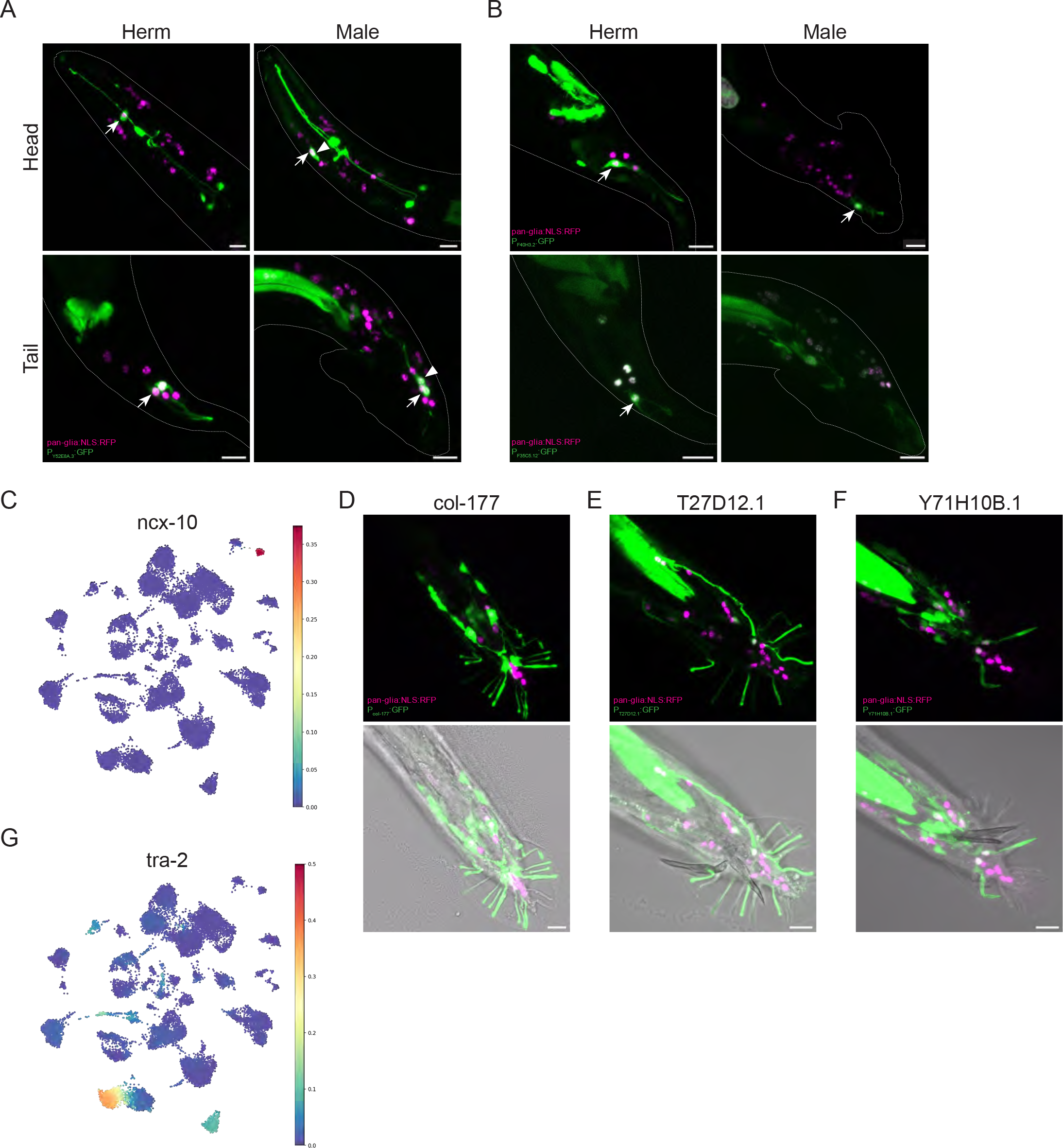
Transcriptional reporters reveal sexually dimorphic expression. (**A**) Z-stack projections of adult hermaphrodite and male heads and tails with dotted outlines showing expression of transcriptional reporter for *Y52E8A*.*3*. In hermaphrodites, expression is observed in AMso glia and Phso1 glia (arrows), as well as pharyngeal neurons. In males, expression is observed in Amso glia (arrow), MCM neuron (arrowhead) in the head and PHD neuron in the tail (arrowhead). Arrow depicts potential Phso1 glia still present in male. (**B**) Top: Z-stack projections of adult hermaphrodite and male tails with dotted outlines showing expression of transcriptional reporter *F40H3*.*2*. In both sexes, expression is observed in Phso2 glia (arrows). Bottom: Z-stack projections of adult hermaphrodite and male tails with dotted outlines showing expression of transcriptional reporter *F35C5.12*. Expression is observed in Phso2 glia (arrow) only in hermaphrodites. (**C**) Glial only batch corrected UMAP showing lack of *ncx-10* gene expression within the glial clusters. (**D**) Z-stack projections of adult male tail showing expression of *col-177* in all ray glia. © Z-stack projections of adult male tail showing expression of *T27D12.1* in a subset of ray glia (specifically in RnSt 1, 3, 5, 7). (**F**) Z-stack projections of adult male tail showing expression of *Y71H10B.1* specifically in RnSt 6/ray 6 glia. (**G**) Glial only batch corrected UMAP showing *tra-2* gene expression within Olso clusters only. Genotypes: Figure 4A: *him-5; PY52E8A.3:GFP, Punc-122:GFP; PmiR-228:nls:RFP, Punc-122:RFP*. Figure 4B: *him- 5; PF40H3.2:GFP, Punc-122:GFP; PmiR-228:nls:RFP, Punc-122:RFP* & *him-5; PF35C5.12:GFP, Punc-122:GFP; PmiR-228:nls:RFP, Punc-122:RFP*. Figure 4D: *him-5; Pcol-177:GFP, Punc-122:GFP; PmiR-228:nls:RFP, Punc-122:RF*P. Figure 4E: *him-5; PT27D12.1:GFP, Punc-122:GFP; PmiR-228:nls:RFP, Punc-122:RFP*. Figure 4F: *him-5; PY71H10B.1:GFP, Punc-122:GFP; PmiR-228:nls:RFP, Punc-122:RFP*. All scale bars = 10μM.

The sole hermaphrodite-specific cluster defines PHso2 glia. This is interesting because while PHso2 is found in both sexes, it is anatomically distinct between the two. Thus, our results indicate that while present in both sexes, PHso2 is a molecularly distinct glial cell in the hermaphrodite. Our studies reveal intriguing complexity. We examined two transcriptional reporters for identified for this cluster *F40H3.2* and *F35C5.12* (Figure S4B). As expected, both genes express in the hermaphrodite PHso2 glia (Figure 4B) However, *F40H3.2* is also expressed in male PHso2 while *F35C5.12* does not (Figure 4B). Thus, although the PHso2 cluster is composed of >95% hermaphrodite cells, it likely has both sex-shared and sex-divergent molecular modules (Figure S2F, cluster 0). We did not uncover a male specific PHso2 cluster, leading us to speculate that either we did not enrich for the male PHso2 during dissociation, or its identity is one of the remaining unidentified male specific cluster.

Anatomically, males have seven types of sex-specific glia clusters, but our clustering reveals there are nine. Thus, at least two additional male glia exhibit dimorphism. We identified one as the PHsh glia. Briefly, the AMsh/PHsh glial cluster (Cluster 27), contains cells from both sexes, and validation of UEGs from this cluster show expression in both sexes and AMsh/PHsh glia (Figure 2F, 2G). However, the male-specific Cluster 10 also maps to PHsh glia, suggesting that its molecular profile is sufficiently distinct to render it to cluster separately. Most genes expressed in Cluster 27 are expressed at lower levels in AMsh/PHsh Cluster 10, rendering differences hard to parse. However, pairwise comparison analysis between the two clusters revealed *ncx-10* as a Cluster 10-specific UEG (Figure 4C). We also note that many UEGs in Cluster 10 are also present in other male-specific clusters and absent in sex-shared clusters, suggesting that these may reveal a male glia-specific signature (i.e. *C17G10.10, F47E1.4* and *W04G3.10,* Figure S4C).

**R**ay **st**ructural cells (RnSt) are the most abundant male-specific glia (9 bilateral, 18 total) that are proposed to have both sheath and socket functions (Lints and Hall, 2005). Surprisingly, our validations did not uncover any clusters specific to these glia. Two possibilities may explain their absence: either the dissociation methods precluded enrichment of these epithelia/hypodermis- embedded cells, or (b) they are related to other tissues and erroneously excluded in our “glia- enrichment” analysis. We did uncover genes that express in RnSt, besides other glia. For example, the pan-sheath marker *kcc-3* expresses in all RnSt (Figure 3D). Likewise, while *col-177* is only expressed in the ILso and OLQso glia in the head region of both males and hermaphrodites, it is also expressed in all the RnSts in males (Figure 4D). Finally, we have uncovered molecular expression profiles of other genes tested in subset of RnSt. For example, *T27D12.1* is enriched in some ILsh/OLsh, and also specifically in RnSt 1, 3, 5, 7 (Figure 4E). Similarly, *Y71H10b.1* is expressed in CEPsh/so glia in both sexes, and only in RnSt 6 in the male (Figure S2J, Figure 4F). This is intriguing because ray 6, which includes RnSt6 and its associated neurons, is anatomically unique in structure compared to other rays (Lints and Hall, 2005). Thus, our data reveal interesting molecular heterogeneity between the different male tail ray cells, which would be a promising area for additional functional investigation. Data for all RnSt expressions are summarized in Table S1.

Finally, we note that Cluster 9 and 12 (OLso), which are sex-shared, express the feminizing gene *tra-2* specifically in the male cell datasets (Figure 4G). Prior gene enrichment analyses also suggest *tra-2* in OLso-associated OLL neurons (Smith et al., 2010). None of the other sex-determination genes (*her-1, fem-1/2/3, tra-1)* have expression in specific glia (Figure S4D). This suggests a specific *tra-2* dependent mechanism for this glia to “feminize” OLso to match molecular profiles of hermaphrodite OLso.

### Global analysis reveals that glia lack canonical DCV release mechanisms

When evaluating our datasets to identify glial clusters (Figure 2E) from neuron and anatomical/epithelia clusters, we were surprised to serendipitously find that *unc-31/CAPS* was not expressed in glial clusters (Figure 5A). This tracks prior reporter studies showing that UNC- 31 is expressed in neurons and vulval muscles (Ailion et al., 1999; Speese et al., 2007). We reconfirmed this by examining *unc-31* GFP transcriptional reporter expression in the pan-glia P*miR-228:NLS* RFP strain. As expected, there was no expression in glia of either sex, but we did see expression emerge in cells in males that retained P*miR-228:*NLS:RFP positive – the presumptive MCM and PHD neurons of the head and tail, respectively (Figure 5B). As UNC- 31/CAPS is required for Ca^2+^-dependent secretion of DCV (Ailion et al., 1999; Speese et al., 2007; Taghert and Veenstra, 2003), this led us to examine DCV biology more closely in glia.

**Figure 5.**
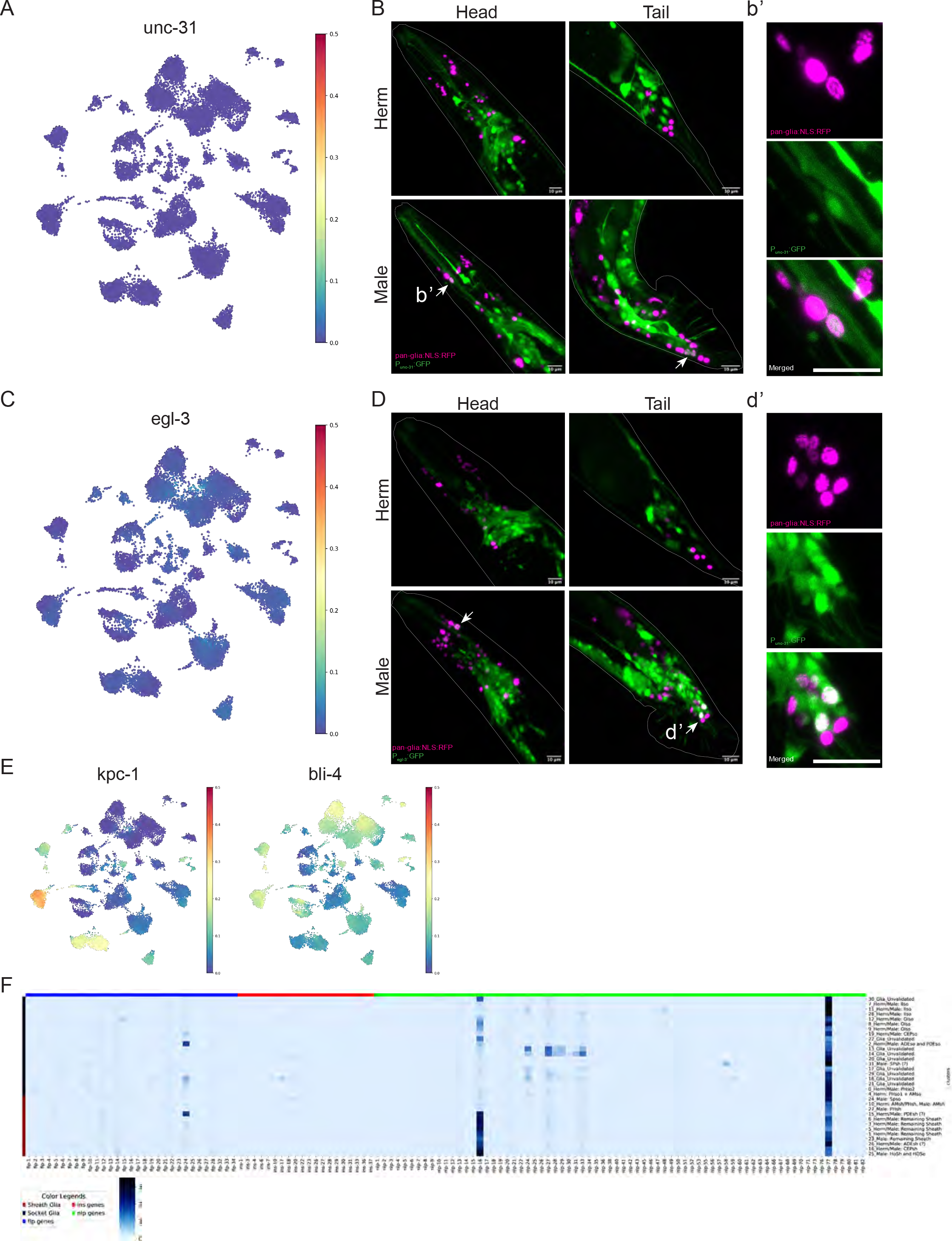
Global analysis reveals that glia lack canonical DCV release and neuropeptide processing mechanisms. (**A**) Glial only batch corrected UMAP showing lack of *unc-31* gene expression in glial clusters. (**B**) Z-stack projections of adult hermaphrodite and male heads and tails with dotted outlines showing expression of transcriptional reporter for *unc-31*. In hermaphrodites, no colocalization is observed. In males, expression is observed in MCM and PHD neurons (arrows). (**b’**) High magnification images and split panels showing expression of *unc-31* in the MCM neuron (lower), but not the AMso glia directly above it. (**C**) Glial only batch corrected UMAP showing lack of *egl-3* gene expression in glial clusters. (**D**) Z-stack projections of adult hermaphrodite and male heads and tails with dotted outlines showing expression of transcriptional reporter for *egl-3*. In hermaphrodites, no colocalization is observed. In males, expression is observed in MCM and PHD neurons (arrows). (**d’**) High magnification images and split panels showing expression of *egl-3* in the PHD neurons. (**E**) Glial only batch corrected UMAP showing *kpc-1* and *bli-4* gene expression in glial clusters. Gene *kpc-1* is especially enriched in the IL/OLso clusters. (**F**) Heatamp showing expression of f*lp* (blue), *ins* (red), and *nlp* (green) neuropeptide genes in the batch corrected glial clusters. Clusters are represented on the y-axis and socket genes are on top (black) while sheath genes are on the bottom (brown). Genotypes: Figure 5B: *him-5; Punc-31:GFP; PmiR-228:nls:RFP, Punc-122:RFP*. Figure 5D: *him-5; Pegl-3:GFP; PmiR-228:nls:RFP, Punc-122:RFP*. All scale bars = 10μM.

Briefly, there are multiple neuropeptide family genes, organized in complex networks in *C. elegans* (Frooninckx et al., 2012; Li et al., 1999; Nathoo et al., 2001; Ripoll-Sánchez et al., 2022). Neuropeptide precursors are first processed by proprotein convertases (mainly *egl-3* but also *kpc-1, bli-4, aex-5* in *C. elegans*). These are then edited by carboxypeptidases (mainly *egl- 21* but also *cpd-1, cpd-2* in *C. elegans)* before being amidated (*pamn-1, pghm-1, pgal-1*) and packaged for release (Hobert, 2013; Van Bael et al., 2018). Neuropeptide signal termination occurs via degradation by different enzyme classes (e.g. *tpp-2, dpt-1, acn-1, dpf-1/2/3/4/6, neprilysins)* (Hobert, 2013). We examined the expression profile of all these genes in our datasets. First, we found that like *unc-31*, the major convertase *egl-3* is not in P*miR-228:NLS* RFP cells, except the male-specific MCM and PHD neurons (Figure 5C, D). This is consistent with prior findings that *egl-3* is exclusively expressed in neurons [Kass]. Hence, *miR-228*-expressing glia do not use either UNC-31 or EGL-3 for DCV biogenesis and exocytosis.

However, we did find expression of transcripts for two of the other convertases (*kpc-1, bli-4*) in glia (Figure 5E). We also found expression of the carboxypeptidase *cpd-1* in glia, and limited expression of all amidation enzymes (*pamn-1, pghm-1, pgal-1*) (Figure S5A). We also found evidence of expression of ***F****MRF-**l**ike **p**eptides* (*flps)*, **ins**ulin-like genes (ins), and ***n****europeptide- **l**ike **p**roteins* (*nlps*) within glial clusters (Figure 5F). Finally, we note varied expression of multiple degradation enzymes (*tpp-2, dpf-1, dpf-2, acn-1*) (Figure S5B). Taken together, we posit that while glia may express some neuropeptides, they process and release DCVs using mechanisms distinct from neurons. How DCVs dock and release in glia in the absence of UNC- 31/CAPS will be an interesting avenue for future research.

## DISCUSSION

In this study we anatomically and molecularly characterized adult *C. elegans* glia encompassing both hermaphrodites and males. We performed snRNA-seq on neuroepithelial glia labeled by pan-glial marker *miR-228*. Adult hermaphrodites contain 50 neuroepithelial glia and males contain an additional 36. Although there was variability in the number of cells per cluster, our final dataset contained 31,410 total cells, giving us >200x coverage on average. This validated dataset of adult glial gene expression provides a resource for glial specific expression in *C. elegans* adults and across sexes.

### Molecular profile of glia across a multicellular nervous system by sex

Prior studies in *C. elegans* have performed either pan-cellular scRNAseq across developmental ages (Cao et al., 2017; Packer et al., 2019; Roux et al., 2022), or neuron-specific scRNAseq at L4 larval development in hermaphrodites (CenGen) (Ripoll-Sánchez et al., 2022)t. Here, we report the complete snRNAseq of adult *C. elegans* glia, across both sexes. Our work complements efforts to profile glia across species (Lago-Baldaia et al., 2022; Özel et al., 2021; Zhang et al., 2014), with the unique benefit that the invariant developmental lineages and glia-neuron contacts in our experimental model, *C. elegans,* allows for single-cell specific validation of our datasets at unprecedented resolution. We further performed machine learning and iterative computational modeling to assign anatomical correlates to all UMAP clusters identified, and validated most clusters functionally *in vivo.* These reveal extensive and variable diversity and dimorphism in glia in this animal.

### Pan-glia and subclass signatures for glia

There is currently no pan-glial-specific marker in *C. elegans.* We used machine-learning models to identify one, which revealed that at least 3 genes are required to reliable call a cluster “Glia” (*brp-1, col-34* and *col-103*). While *brp-1* alone expresses broadly across glia, we have not been able to verify its expression *in vivo.* This may explain why prior studies have not revealed any single such genetic marker. All genes are evolutionarily conserved, so it will be interesting to evaluate if this pan-glia signature profile is broadly conserved in other species. The validation of these genes using an independently published atlas of all cells in *C. elegans* during various stages of adulthood demonstrated the consistency of the pan-glial signatures across animal age and that our glial signature profiling can accurately identify glia in independently obtained snRNA-seq datasets (Roux et al., 2022).

Our modeling also revealed markers for the two anatomical sub-classes of glia, sheath (*kcc-3, ttr- 43)* and socket (*zipt-2.2, cnc-10)*. That there are no broadly shared sheath/socket markers identified is interesting and indicates that these mark the terminal hierarchical designation of two entirely distinct glia types, rather than subsets of a basal glia-state. This is distinct from different neuron classes still sharing a “pan-neuron” signature profile.

Curiously, molecular profiles of some male-specific glia (i.e SPso/sh, PCso/sh, HOsh/so) unexpectedly did not track sheath/socket predictions based on anatomical identity. We suggest that these may allow future evaluation of sheath versus socket functions, and these glia may express other factors that enable “anatomical” identity of one subclass, while molecular profile of another. Why this is observed only for male-specific glia will also be interesting to assess.

These studies also reveal variable heterogeneity within glial anatomical classes. For example, prior transcriptomic profiling had shown that although the ILso glia are produced by three symmetric pairs of lineages, the developmental trajectories formed by the ILso progenitors and their terminal descendants were discontinuous in UMAP space, implying disparate transcriptional profiles (Packer et al., 2019). In line with this observation, we also note heterogeneity within the IL/OLso clusters. In contrast, the IL/OLsh glia, although non-identical, are transcriptionally significantly similar.

### Glial sex dimorphism

The anatomical differences between the glia of the two sexes are well-known, but the relationship between anatomy and glial molecular identity for sex-specific or sex-shared glia has not been explored before. Our dataset uncovered sex-specific and sex-shared clusters.

Interestingly, although the AMso and PHso1 of the two sexes are anatomically distinct, and have different developmental fates in the larval stage, they contain the same transcriptional profile.

Moreover, some glial cells are present in both sexes but are molecularly distinct in each sex (i.e. PHsh and PHso2). Further, we note that there is non-uniformity in sex dimorphism across different glia, and may arise from distinct glia-specific mechanisms. Thus, the molecular diversity and dimorphism across glia extends beyond anatomy. The data presented here provide molecular tools to examine this further. We speculate that these molecular differences may be related to functional differences that have not yet been explored.

### Glia and neurons process neuropeptides through different machinery

Neuropeptides are the most diverse class of signaling molecules in the brain (Burbach, 2011). Even though there is evidence of expression and release of neuropeptides in all types of mammalian glia, little is known about neuropeptide storage and release mechanisms (Ubink et al., 2003). Our data suggests that the protein UNC-31/CAPS, which is required for Ca2+- dependent secretion of DCV in neurons, is not expressed in glia in *C. elegans*. Similarly, the proprotein convertase *egl-3* that is expressed in the nervous system is also not expressed in glia. Interestingly, others have shown that rescue of *unc-31* and *egl-3* in CEPsh glia can fully rescues lifespan extension and lead to an increase in activation of the UPR^ER^ in the intestine (Frakes et al., 2020). This suggests that *unc-31* and *egl-3* can compensate for the machinery required for processing and releasing peptides in glia. While further examining DCV biology components, we did find expression of two other convertases (*kpc-1, bli-4*) and carboxypeptidase *cpd-1* in glia.

Because we found evidence of expression of several neural peptides within distinct glial clusters, further investigation of the mechanisms by which glia process and release DCVs, particularly in the absence of UNC-31/CAPS is important.

Lastly, we acknowledge that our study has some limitations. Because of 3′ bias, our data does not reveal which specific isoform is expressed in each cell. Low abundance transcripts may be underrepresented in snRNA-seq data, particularly in clusters with relatively few cells.

Additionally, manual characterization of clusters is prone to error since marker genes are often expressed in multiple clusters and correspond to multiple cell type.

Overall, this molecular atlas, deposited and searchable at www.wormglia.org provides comprehensive information on the diversity and differences between male and hermaphrodite glial cells throughout the nervous system of an adult organism.

## ACKNOWLEDGEMENTS

We thank the Singhvi and Setty labs for discussions and comments on the manuscript and Robert Waterston and Chau Huynh for early discussions on dissociation protocols. This work was funded by a Washington Research Foundation Postdoctoral award to MP; NIH/NIGMS R35GM147125 to MS; and Simons Foundation/SFARI grant (488574), Esther A. & Joseph Klingenstein Fund and NIH/NINDS funding (NS114222) to AS. This work was performed while AS was a Glenn Foundation for Medical Research and AFAR Junior Faculty Grant Awardee. AS also sincerely thanks philanthropic supporters to her laboratory. Some work was performed at the Fred Hutch Shared Resources Core Facilities. We sincerely apologize if we missed citing works due to our oversight or space considerations.

## AUTHOR CONTRIBUTIONS

MP, AS and MS conceptualized all aspects of this study. MP performed all snRNA-seq experiments and validations with RSM, NT, and LS assisting on transgenic reporter constructions. VS assisted in sorting male animals. MP wrote the manuscript with AS. EQ and MS contributed to writing the methods. EQ performed the computational analytics under primary supervision from MS. MP, AS, MS and EQ analyzed all results.

## REFERENCES

1. Ailion, M., Inoue, T., Weaver, C.I., Holdcraft, R.W., Thomas, J.H., 1999. Neurosecretory control of aging in *Caenorhabditis elegans*. Proc. Natl. Acad. Sci. 96, 7394–7397. https://doi.org/10.1073/pnas.96.13.7394

2. Altun, Z.F., Hall, D.H., 2003. WormAtlas Hermaphrodite Handbook - Nervous System - Neuronal Support Cells. WormAtlas. https://doi.org/10.3908/wormatlas.1.19

3. Bacaj, T., Tevlin, M., Lu, Y., Shaham, S., 2008. Glia Are Essential for Sensory Organ Function in *C. elegans*. Science 322, 744–747. https://doi.org/10.1126/science.1163074

4. Burbach, J.P.H., 2011. What Are Neuropeptides?, in: Merighi, A. (Ed.), Neuropeptides, Methods in Molecular Biology. Humana Press, Totowa, NJ, pp. 1–36. https://doi.org/10.1007/978-1-61779-310-3_1

5. Cao, J., Packer, J.S., Ramani, V., Cusanovich, D.A., Huynh, C., Daza, R., Qiu, X., Lee, C., Furlan, S.N., Steemers, F.J., Adey, A., Waterston, R.H., Trapnell, C., Shendure, J., 2017. Comprehensive single-cell transcriptional profiling of a multicellular organism. Science 357, 661–667. https://doi.org/10.1126/science.aam8940

6. Chasnov, J.R., Chow, K.L., 2002. Why Are There Males in the Hermaphroditic Species *Caenorhabditis elegans* ? Genetics 160, 983–994. https://doi.org/10.1093/genetics/160.3.983

7. Cook, S.J., Jarrell, T.A., Brittin, C.A., Wang, Y., Bloniarz, A.E., Yakovlev, M.A., Nguyen, K.C.Q., Tang, L.T.-H., Bayer, E.A., Duerr, J.S., Bülow, H.E., Hobert, O., Hall, D.H., Emmons, S.W., 2019. Whole-animal connectomes of both Caenorhabditis elegans sexes. Nature 571, 63–71. https://doi.org/10.1038/s41586-019-1352-7

8. Crowley, L.C., Scott, A.P., Marfell, B.J., Boughaba, J.A., Chojnowski, G., Waterhouse, N.J., 2016. Measuring Cell Death by Propidium Iodide Uptake and Flow Cytometry. Cold Spring Harb. Protoc. 2016, pdb.prot087163. https://doi.org/10.1101/pdb.prot087163

9. Frakes, A.E., Metcalf, M.G., Tronnes, S.U., Bar-Ziv, R., Durieux, J., Gildea, H.K., Kandahari, N., Monshietehadi, S., Dillin, A., 2020. Four glial cells regulate ER stress resistance and longevity via neuropeptide signaling in *C. elegans*. Science 367, 436–440. https://doi.org/10.1126/science.aaz6896

10. Frooninckx, L., Van Rompay, L., Temmerman, L., Van Sinay, E., Beets, I., Janssen, T., Husson, S.J., Schoofs, L., 2012. Neuropeptide GPCRs in C. elegans. Front. Endocrinol. 3. https://doi.org/10.3389/fendo.2012.00167

11. Fung, W., Wexler, L., Heiman, M.G., 2020. Cell-type-specific promoters for *C. elegans* glia. J. Neurogenet. 34, 335–346. https://doi.org/10.1080/01677063.2020.1781851

12. Hall, D.H., 1977. The posterior nervous system of the nematode Caenorhabditis elegans. California Institute of Technology.

13. Herculano-Houzel, S., Dos Santos, S., 2018. You Do Not Mess with the Glia. Neuroglia 1, 193–219. https://doi.org/10.3390/neuroglia1010014

14. Hobert, O., 2013. The neuronal genome of Caenorhabditis elegans. WormBook 1–106. https://doi.org/10.1895/wormbook.1.161.1

15. Hobert, O., 2010. Neurogenesis in the nematode Caenorhabditis elegans. WormBook. https://doi.org/10.1895/wormbook.1.12.2

16. Kaletsky, R., Lakhina, V., Arey, R., Williams, A., Landis, J., Ashraf, J., Murphy, C.T., 2016. The C. elegans adult neuronal IIS/FOXO transcriptome reveals adult phenotype regulators. Nature 529, 92–96. https://doi.org/10.1038/nature16483

17. Katz, M., Corson, F., Iwanir, S., Biron, D., Shaham, S., 2018. Glia Modulate a Neuronal Circuit for Locomotion Suppression during Sleep in C. elegans. Cell Rep. 22, 2575–2583. https://doi.org/10.1016/j.celrep.2018.02.036

18. Korsunsky, I., Millard, N., Fan, J., Slowikowski, K., Zhang, F., Wei, K., Baglaenko, Y., Brenner, M., Loh, P., Raychaudhuri, S., 2019. Fast, sensitive and accurate integration of single-cell data with Harmony. Nat. Methods 16, 1289–1296. https://doi.org/10.1038/s41592-019-0619-0

19. Lago-Baldaia, I., Cooper, M., Seroka, A., Trivedi, C., Powell, G.T., Wilson, S., Ackerman, S., Fernandes, V.M., 2022. A *Drosophila* glial cell atlas reveals that transcriptionally defined cell types can be morphologically diverse (preprint). Neuroscience. https://doi.org/10.1101/2022.08.01.502305

20. Li, C., Kim, K., Nelson, L.S., 1999. FMRFamide-related neuropeptide gene family in Caenorhabditis elegans. Brain Res. 848, 26–34. https://doi.org/10.1016/S0006-8993(99)01972-1

21. Lints, R., Hall, D.H., 2005. WormAtlas Male Handbook - Neuronal Support Cells - Rays. WormAtlas. https://doi.org/10.3908/wormatlas.2.10

22. Malin, J.A., Kinet, M.J., Abraham, M.C., Blum, E.S., Shaham, S., 2016. Transcriptional control of non-apoptotic developmental cell death in C. elegans. Cell Death Differ. 23, 1985– 1994. https://doi.org/10.1038/cdd.2016.77

23. McInnes, L., Healy, J., Saul, N., Großberger, L., 2018. UMAP: Uniform Manifold Approximation and Projection. J. Open Source Softw. 3, 861. https://doi.org/10.21105/joss.00861

24. Molina-García, L., Lloret-Fernández, C., Cook, S.J., Kim, B., Bonnington, R.C., Sammut, M., O’Shea, J.M., Gilbert, S.P., Elliott, D.J., Hall, D.H., Emmons, S.W., Barrios, A., Poole, R.J., 2020. Direct glia-to-neuron transdifferentiation gives rise to a pair of male-specific neurons that ensure nimble male mating. eLife 9, e48361. https://doi.org/10.7554/eLife.48361

25. Nathoo, A.N., Moeller, R.A., Westlund, B.A., Hart, A.C., 2001. Identification of *neuropeptide-like protein* gene families in *Caenorhabditis elegans* and other species. Proc. Natl. Acad. Sci. 98, 14000–14005. https://doi.org/10.1073/pnas.241231298

26. Nguon, K., Ladd, B., Baxter, M.G., Sajdel-Sulkowska, E.M., 2005. Sexual dimorphism in cerebellar structure, function, and response to environmental perturbations, in: Progress in Brain Research. Elsevier, pp. 341–351. https://doi.org/10.1016/S0079-6123(04)48027-3

27. Özel, M.N., Simon, F., Jafari, S., Holguera, I., Chen, Y.-C., Benhra, N., El-Danaf, R.N., Kapuralin, K., Malin, J.A., Konstantinides, N., Desplan, C., 2021. Neuronal diversity and convergence in a visual system developmental atlas. Nature 589, 88–95. https://doi.org/10.1038/s41586-020-2879-3

28. Packer, J.S., Zhu, Q., Huynh, C., Sivaramakrishnan, P., Preston, E., Dueck, H., Stefanik, D., Tan, K., Trapnell, C., Kim, J., Waterston, R.H., Murray, J.I., 2019. A lineage-resolved molecular atlas of *C. elegans* embryogenesis at single-cell resolution. Science 365, eaax1971. https://doi.org/10.1126/science.aax1971

29. Pedregosa, F., Varoquaux, G., Gramfort, A., Michel, V., Thirion, B., Grisel, O., Blondel, M., Müller, A., Nothman, J., Louppe, G., Prettenhofer, P., Weiss, R., Dubourg, V., Vanderplas, J., Passos, A., Cournapeau, D., Brucher, M., Perrot, M., Duchesnay, É., 2012. Scikit-learn: Machine Learning in Python. https://doi.org/10.48550/ARXIV.1201.0490

30. Ripoll-Sánchez, L., Watteyne, J., Sun, H., Fernandez, R., Taylor, S.R., Weinreb, A., Hammarlund, M., Miller, D.M., Hobert, O., Beets, I., Vértes, P.E., Schafer, W.R., 2022. The neuropeptidergic connectome of *C. elegans* (preprint). Neuroscience. https://doi.org/10.1101/2022.10.30.514396

31. Roux, A.E., Yuan, H., Podshivalova, K., Hendrickson, D., Kerr, R., Kenyon, C., Kelley, D.R., 2022. The complete cell atlas of an aging multicellular organism (preprint). Genomics. https://doi.org/10.1101/2022.06.15.496201

32. Sammut, M., Cook, S.J., Nguyen, K.C.Q., Felton, T., Hall, D.H., Emmons, S.W., Poole, R.J., Barrios, A., 2015. Glia-derived neurons are required for sex-specific learning in C. elegans. Nature 526, 385–390. https://doi.org/10.1038/nature15700

33. Schwarz, J.M., Bilbo, S.D., 2012. Sex, glia, and development: Interactions in health and disease. Horm. Behav. 62, 243–253. https://doi.org/10.1016/j.yhbeh.2012.02.018

34. Setty, M., Kiseliovas, V., Levine, J., Gayoso, A., Mazutis, L., Pe’er, D., 2019. Characterization of cell fate probabilities in single-cell data with Palantir. Nat. Biotechnol. 37, 451–460. https://doi.org/10.1038/s41587-019-0068-4

35. Shaham, S., 2010. Chemosensory organs as models of neuronal synapses. Nat. Rev. Neurosci. 11, 212–217. https://doi.org/10.1038/nrn2740

36. Simerly, R.B., 2002. Wired for Reproduction: Organization and Development of Sexually Dimorphic Circuits in the Mammalian Forebrain. Annu. Rev. Neurosci. 25, 507–536. https://doi.org/10.1146/annurev.neuro.25.112701.142745

37. Singhvi, A., Liu, B., Friedman, C.J., Fong, J., Lu, Y., Huang, X.-Y., Shaham, S., 2016. A Glial K/Cl Transporter Controls Neuronal Receptive Ending Shape by Chloride Inhibition of an rGC. Cell 165, 936–948. https://doi.org/10.1016/j.cell.2016.03.026

38. Singhvi, A., Shaham, S., 2019. Glia-Neuron Interactions in *Caenorhabditis elegans*. Annu. Rev. Neurosci. 42, 149–168. https://doi.org/10.1146/annurev-neuro-070918-050314

39. Smith, C.J., Watson, J.D., Spencer, W.C., O’Brien, T., Cha, B., Albeg, A., Treinin, M., Miller, D.M., 2010. Time-lapse imaging and cell-specific expression profiling reveal dynamic branching and molecular determinants of a multi-dendritic nociceptor in C. elegans. Dev. Biol. 345, 18–33. https://doi.org/10.1016/j.ydbio.2010.05.502

40. Soreq, L., Rose, J., Soreq, E., Hardy, J., Trabzuni, D., Cookson, M.R., Smith, C., Ryten, M., Patani, R., Ule, J., 2017. Major Shifts in Glial Regional Identity Are a Transcriptional Hallmark of Human Brain Aging. Cell Rep. 18, 557–570. https://doi.org/10.1016/j.celrep.2016.12.011

41. Speese, S., Petrie, M., Schuske, K., Ailion, M., Ann, K., Iwasaki, K., Jorgensen, E.M., Martin, T.F.J., 2007. UNC-31 (CAPS) Is Required for Dense-Core Vesicle But Not Synaptic Vesicle Exocytosis in Caenorhabditis elegans. J. Neurosci. 27, 6150–6162. https://doi.org/10.1523/JNEUROSCI.1466-07.2007

42. Stout Jr., R.F., Verkhratsky, A., Parpura, V., 2014. Caenorhabditis elegans glia modulate neuronal activity and behavior. Front. Cell. Neurosci. 8. https://doi.org/10.3389/fncel.2014.00067

43. Sulston, J.E., Albertson, D.G., Thomson, J.N., 1980. The Caenorhabditis elegans male: Postembryonic development of nongonadal structures. Dev. Biol. 78, 542–576. https://doi.org/10.1016/0012-1606(80)90352-8

44. Sulston, J.E., Horvitz, H.R., 1977. Post-embryonic cell lineages of the nematode, Caenorhabditis elegans. Dev. Biol. 56, 110–156. https://doi.org/10.1016/0012-1606(77)90158-0

45. Taghert, P.H., Veenstra, J.A., 2003. Drosophila Neuropeptide Signaling, in: Advances in Genetics. Elsevier, pp. 1–65. https://doi.org/10.1016/S0065-2660(03)01001-0

46. Taylor, S.R., Santpere, G., Weinreb, A., Barrett, A., Reilly, M.B., Xu, C., Varol, E., Oikonomou, P., Glenwinkel, L., McWhirter, R., Poff, A., Basavaraju, M., Rafi, I., Yemini, E., Cook, S.J., Abrams, A., Vidal, B., Cros, C., Tavazoie, S., Sestan, N., Hammarlund, M., Hobert, O., Miller, D.M., 2021. Molecular topography of an entire nervous system. Cell 184, 4329–4347.e23. https://doi.org/10.1016/j.cell.2021.06.023

47. Traag, V.A., Waltman, L., van Eck, N.J., 2019. From Louvain to Leiden: guaranteeing well-connected communities. Sci. Rep. 9, 5233. https://doi.org/10.1038/s41598-019-41695-z

48. Ubink, R., Calza, L., Hökfelt, T., 2003. ‘Neuro’-peptides in glia: Focus on NPY and galanin. Trends Neurosci. 26, 604–609. https://doi.org/10.1016/j.tins.2003.09.003

49. Van Bael, S., Watteyne, J., Boonen, K., De Haes, W., Menschaert, G., Ringstad, N., Horvitz, H.R., Schoofs, L., Husson, S.J., Temmerman, L., 2018. Mass spectrometric evidence for neuropeptide-amidating enzymes in. J. Biol. Chem. 293, 6052–6063. https://doi.org/10.1074/jbc.RA117.000731

50. van Dijk, D., Sharma, R., Nainys, J., Yim, K., Kathail, P., Carr, A.J., Burdziak, C., Moon, K.R., Chaffer, C.L., Pattabiraman, D., Bierie, B., Mazutis, L., Wolf, G., Krishnaswamy, S., Pe’er, D., 2018. Recovering Gene Interactions from Single-Cell Data Using Data Diffusion. Cell 174, 716–729.e27. https://doi.org/10.1016/j.cell.2018.05.061

51. Wallace, S.W., Singhvi, A., Liang, Y., Lu, Y., Shaham, S., 2016. PROS-1/Prospero Is a Major Regulator of the Glia-Specific Secretome Controlling Sensory-Neuron Shape and Function in C. elegans. Cell Rep. 15, 550–562. https://doi.org/10.1016/j.celrep.2016.03.051

52. Ward, S., Thomson, N., White, J.G., Brenner, S., 1975. Electron microscopical reconstruction of the anterior sensory anatomy of the nematodecaenorhabditis elegans. J. Comp. Neurol. 160, 313–337. https://doi.org/10.1002/cne.901600305

53. Wolf, F.A., Angerer, P., Theis, F.J., 2018. SCANPY: large-scale single-cell gene expression data analysis. Genome Biol. 19, 15. https://doi.org/10.1186/s13059-017-1382-0

54. Yoshida, A., Nakano, S., Suzuki, T., Ihara, K., Higashiyama, T., Mori, I., 2016. A glial K ^+^ /Cl ^−^ cotransporter modifies temperature-evoked dynamics in *Caenorhabditis elegans* sensory neurons. Genes Brain Behav. 15, 429–440. https://doi.org/10.1111/gbb.12260

55. Zhang, Y., Barres, B.A., 2010. Astrocyte heterogeneity: an underappreciated topic in neurobiology. Curr. Opin. Neurobiol. 20, 588–594. https://doi.org/10.1016/j.conb.2010.06.005

56. Zhang, Y., Chen, K., Sloan, S.A., Bennett, M.L., Scholze, A.R., O’Keeffe, S., Phatnani, H.P., Guarnieri, P., Caneda, C., Ruderisch, N., Deng, S., Liddelow, S.A., Zhang, C., Daneman, R., Maniatis, T., Barres, B.A., Wu, J.Q., 2014. An RNA-Sequencing Transcriptome and Splicing Database of Glia, Neurons, and Vascular Cells of the Cerebral Cortex. J. Neurosci. 34, 11929–11947. https://doi.org/10.1523/JNEUROSCI.1860-14.2014

