## Supplemental Materials and Figures for "Molecular heterogeneity of *C. elegans* glia across sexes"

This pdf includes following supplemental materials:

Materials and Methods

Supplemental Figures S1-S5

Supplemental Figure Legends S1-S5

Reagents

Supplemental References

*To whom correspondence should be addressed:

#### EXPERIMENTAL PROCEDURES

##### *C. elegans* methods

*C. elegans* were cultured and maintained at 20ºC on OP50 plates, as previously described (Brenner, 1974; Stiernagle, 2006).

##### Strains

Strains were sourced from the CGC, funded by NIH Office of Research Infrastructure Programs (P40 OD010440).

##### Preparation of adults for dissociation

Worms were grown on 10 cm, standard 1% NGM agar plates seeded with *E.coli* strain OP50. To obtain synchronized day 1 adult worms, we first obtained embryos by performing hypochlorite treatment (1M NaOH + 4% NaOCl freshly made) on adult hermaphrodites. The embryos hatched overnight (16-20 hours) on unseeded NGM agar plates and starved L1 arrested worms were recovered on 10 cm OP50-seeded plates at 20C for 64-68 hours. For hermaphrodite selection: For each sample, about 50,000 day 1 adult worms were washed off plates with M9 into 50 ml conical tubes that were fitted with a 50um pluriStrainer® filter to separate day 1 adults from embryos or hatched L1s. The pure day 1 adult hermaphrodite population was transferred to 15 ml conical tubes and further washed 3x with M9 with centrifugation at 1200 rcf for 1 min. For male enrichment: For each sample, about 150,000 day 1 adult worms were washed off plates with M9 into 50 ml conical tubes that were fitted with stacked 50 um, 40 um, and 30 um pluriStrainer® filters to separate adult males from hermaphrodites. The stacked filtration was repeated one more time. The worms from the 30 um filter were mix of about 80-90% males, plus young day 1 adult hermaphrodites. To further enrich for a male only population, we used the COPAS worm sorter to select animals based on size and lack of $P_{mig\text{-}24}$: mVenus staining and recovered 50,000 males. The males were further washed 3x with M9 with centrifugation at 1200 rcf for 1 min.

**Dissociation**

Single nuclei suspensions were obtained as described (Kaletsky et al., 2016) with some modifications to further dissociate the cell membranes. For the hermaphrodite Worms were split into 1.6ml clear tubes with M9, containing 250ul pellets. We used a tabletop mini centrifuge to pellet worms during all the dissociation steps. After removing most of the M9 solution, worms were quickly washed once in 500ul lysis buffer (200mM DTT, 0.25% SDS, 20mM Hepes pH 8.0, 3% sucrose), and then lysed with 750ul lysis buffer for 5 min for hermaphrodites and 3 minutes for males. We shortened the lysis for males because longer lysis time caused the males to rupture during the downstream M9 washing steps. We monitored cuticle disruption by examining 1μL of sample under a microscope. When lysis was complete, worms had a slightly blunt head, but were not ruptured. We stopped the reaction by adding 500ul M9 and then did 6 quick washed in M9. Worms were then dissociated by adding 500 ul freshly made 20mg/ml Pronase (resuspended in water) to the worm pellet for 15-20 minutes depending on the lot of Pronase. Males were dissociated in 15 mg/ml Pronase. During the Pronase incubation, worms were dissociated by pipetting vigorously with a P1000 pipette set to 400ul every 45 seconds, rotating between samples for the first 10 minutes, followed by trituration with a P200 pipette set to 200 ul for remaining 5-10 minutes. Worms were quickly spun down between triturations so that the worm pellet size could be monitored. Between pipetting, 2ul of worms were examined on a dissecting scope to assess level of dissociation. We continued dissociating until large fragments were gone and the pellet of worms was less than 20 ul when quickly centrifuged. We noticed that during this over digestion step, the cell membranes would also digest. To stop the digestion, 750µL of ice-cold 1xPBS/10%FBS was added to the tube. The dissociated nuclei were transferred to a 1 ml syringe using a 27-gauge needle. The needle was replaced by a 5uM syringe filter that was pre-wetted with ice cold PBS. The cells were gently passed through the syringe filter directly into the FACS collection tube on ice. Protector RNAse Inhibitor was added to the following solutions: Pronase, PBS-FBS, and PBS-L15-FBS to get final concentration of 2U/ul inhibitor.

**Cell sorting and single nuclear RNA sequencing**

The nuclei were sorted with a SH800S cell sorter fitted with the 100µm nozzle running at a speed of 10-12k events per second using the PE filter set. For the hermaphrodite samples, we included age matched N2 worms as negative control to set FACS gates. For the males samples, we used *him-5* males as negative controls. Both negative controls were taken through the same protocol as above. Nuclei were sorted into 1:1 1xPBS:L15+30%FBS 2U/µl Protector RNAse inhibitor, transferred to a low bind 2 ml tube and centrifuged 9600xg for 5 min at 4C. Once an RFP+ pellet was confirmed under a fluorescent dissection scope, the nuclei were gently resuspended in 1xPBS10%FBS buffer aiming for a final concentration of 1000 nuclei/uL based on the number of nuclei sorted. The nuclei were then loaded into a hemocytometer and counted to determine the final concentration. We targeted 10,000 cells per sample and used the 10X Chromium platform using 10X NextGem v3.1 chemistry. cDNA was amplified for 12 cycles and the index PCR was cycled 12 times. Final libraries were brought to 10 nM concentration and processed on an Illumina HiSeq2500 sequencer. The males and hermaphrodite samples were prepared and sequenced on different days.

**RT-PCR**

To perform RT-PCR, ~200,000 RFP+ nuclei were sorted directly into Trizol LS and then stored at −80°C until samples were ready for RNA extraction. The whole animal controls consisted of dissociated nuclei of N2 worms that were sitting on ice during FACS to mimic the same amount of time the RFP+ samples were being sorted –100ul of this nuclei suspension was added to Trizol LS after FACS was complete. After thawing, the samples were incubated at 65C for 5 min with vortexing every minutes. One fifth volume of chloroform was added to each sample and mixed in a 5' Prime- Heavy Phase Lock Gel, and samples were then centrifuged (12,000xg for 15 min). The aqueous phase was removed and RNA was isolated using Qiagen RNeasy MinElute Cleanup Kit. RNA was quantified using the Qubit fluorometer. DNAse digestion and RNA reverse transcription was performed using Qiagen QuantiTect Reverse Transcription Kit. The resulting cDNA was diluted 1:5 and 2 ul was used for a single RT-PCR reaction. All real time assays were performed using TaqMan gene expression assays (ThermoFisher) and TaqMan Fast Advanced Master Mix on a QuantStudio™ 5 system (ThermoFisher). Gene *pmp-3* (TaqMan assay Ce02485188_m1) was used as the control housekeeping gene. The following TaqMan assays were also utilized: *unc-119* (TaqMan assay Ce02452615_g1), *F53F4.13* (TaqMan assay Ce02484052_g1), *kcc-3* (TaqMan assay Ce02434964_g1), *ptr-10* (Taqman assay Ce02418075_g1).

**Microscopy, Image Processing and Analysis**

Day 1 animals were immobilized with 40mM sodium azide. Images were collected on a Zeiss 780 LSM NLO with a 40x/1.3NA Plan Neofluar oil-immersion. Image processing was done in FIJI ImageJ.

**Single-cell data analysis**

10X Cellranger (v5.0.1) was used to generate the count matrices for each sample. The 3' untranslated regions (UTRs) in the reference transcriptome (WormBase WS280) have been previously reported to be too short. Therefore, following the previously published approaches, we extended 3' UTRs in the WS280 genome. Specifically, for genes not overlapping with a downstream gene, we extended their 3' UTRs by 50, 100, 150, 200, 250, 300, 400 and 500bp. In case the extended sequence collided with a downstream gene, it was trimmed back to the base next to the gene. The unmodified and eight modified GTF files were each used to build a reference genome for 10X Cellranger. Protein-coding genes and non-coding RNA genes (lincRNAs, ncRNAs and antisense RNAs) were included for analyses. 10X Cellranger was used to generate a count matrix for each sample relative to each reference. The optimal 3' UTR extension was defined as follows: 20 reads were considered as a significant gain by an extension interval. For each gene, the cumulative sums from 3' (the longest extension) to 5' was calculated after subtracting 20 from read increment in each extension interval. The optimal extension for a given sample was set to the point which had the smallest cumulative cum of less than 0. If cumulative sums were all greater than zero, the optimal extension was set to 500bp. The final extension of a gene was set to the point supported by most samples and by at least two samples. Cell-containing droplets by 10X Cellranger were used for downstream analysis (16687 cells in hermaphrodites and 14723 cells in males replicates respectively, for a total of 31410 cells measured across 28045 genes). Housekeeping genes were also excluded from downstream analyses. The four samples were concatenated resulting in a final count matrix of 31410 cells and 12389 genes.

**Data processing**

Data processing was performed using the scanpy package (Wolf et al., 2018) with default parameters unless specified. Raw counts were normalized by dividing the counts by total counts per cell. The normalized data was multiplied by the median of total counts across cells to avoid numerical issues and log-transformed with a pseudo count of 1. Feature selection was then performed to select the top 2500 most highly variable genes (using **scanpy.pp.highly_variable_genes**), which was used as input for principal component analysis with 50 components. The PCs were used as input for generating UMAPs (McInnes et al., 2018) (with **min_dist=0.2**) and clustering with leiden (Traag et al., 2019), resulting in 51 clusters. MAGIC (van Dijk et al., 2018) imputation was used to visualize gene expression on UMAPs.

**Annotation of glial, neuronal, and anatomical compartments**

We utilized a multi-pronged approach to annotate each of the 51 clusters as belonging to one of glial, neuronal, and anatomical compartments. First, we identified differentially expressed genes for each cluster using **scanpy.tl.rank_gene_groups** by comparing gene expression in each cluster to all other cells. of multiple approaches to iteratively profile individual clusters in our data (adjusted **p-value < 1e-3 , logFC > 1.5)**. We performed a literature survey to identify cell-types and compartments where possible (Supplementary table with markers, cell-types, compartments, references including WormBase genes, Supplementary dot plots showing gene expression).

Next, we utilized the CeNGEN dataset (Taylor et al., 2021)to identify neuronal, and other non-glial compartments. Single cell count matrices and annotations were downloaded and data was processed as described in the "Data processing" section. Differential expression was performed to identify gene signatures that define each cell-type (adjusted **p-value < 1e-3**, **logFC > 1.5).** CeNGEN cell-type signatures were used to derived gene signatures for each cell in our dataset: We first z-scored the expression of each gene across all cells and a signature score was computed for each cell and determined as the average of the z-scored expression across the genes that define the signature. Gene signature scores were used to associate compartments to clusters based on manual inspection of the plotted individual scores for a geneset per cluster.

**Batch effect correction between hermaphrodite and male cells**

We excluded mitochondrial genes prior to batch correction. We then identified highly variable genes separately amongst the hermaphrodite and male cells (2500 genes for each sexes) and used the union of these highly variable genes for downstream analysis (3355 genes) as input to PCA (50 components), Harmony (Korsunsky et al., 2019) was to perform batch effect correction and the corrected PCs were used as input for UMAP and leiden clustering. We identified 43 clusters following batch-correction. Cell-type compartments were transferred by computing the fractions of glia, neuronal and anatomical cells within each cluster and assigning labels to each cluster based on the highest proportion of cells that make up a particular cluster. Batch corrected data was used for visualizations and for identification of pan-glial marker genes.

**Glial compartment batch correction**

Count matrix was subset to contain only the annotated glial clusters. Batch correction was performed as described above after reselecting highly variable genes using only the glial cells (3626 genes). Leiden clustering led to identification of 32 clusters. Batch corrected glial cell data was then subsequently used for further identification of sheath and socket marker genes.

#### ANNOTATION OF GLIAL CELLS

**Glial cell-types and markers**

We observed that standard differential expression analysis that compares cells of one cluster to all other cells was insufficient to clearly identify genes that are uniquely expressed in each cluster. Therefore, we devised a pairwise analysis scheme to identify cluster-specific genes and markers for glial cell-types. Using batch corrected glial data, we performed differential expression analysis using **scanpy.tl.rank_gene_groups** to compare cells of a particular cluster with cells of every other cluster separately. We then enumerated the number of comparisons in which a gene was significantly higher (adjusted **p-value < 1e-25, logFC > 1.5** for more stringent filtering criteria to ensure specificity). Genes were ranked based on their differential frequency and visualized using heatmaps. Candidates for *in vivo* validation were selected based on their differential frequency. Finally, clusters are annotated as specific cell-types based on their in vivo anatomical locations and previously characterized marker genes

#### SEX-SPECIFIC AND SHARED GLIAL CELL-TYPES

**Manual Quantification**

We quantified the fractions of male and hermaphrodite samples within a given cluster for each type of dataset (whole data and glia only data) for non-batch corrected and or batch corrected data. Sex specificity labels for each cluster was assigned based on a threshold (whole data threshold = 95%, glia only data = 90%), that is, if a cluster was at least above a certain percentage of one sex or the other it was determined as sex specific for that sex otherwise it was labeled as both/shared. The thresholds were chosen such that after quantification, should reflect the true biology of sex-specific glia. In this case we expected one or two hermaphrodite specific glial clusters to account for hermaphrodite specific glia.

**Determination of pan-glial markers**

We used a supervised classification framework to identify the core set of markers that can accurately define all glial cells. We trained a logistic regression model to distinguish glial cells from non-glial cells using MAGIC imputed gene expression data and devised a feature ranking scheme to identify a small set of markers that span all glial cells. The model was trained using the hermaphrodite cells from the batch corrected data and generalize to male cells The following criteria were used for selection of genes as features: (i) Genes that are highly variable in either hermaphrodite or male cells and (ii) Genes should be detected in at least of 40% of cells in at least one cluster.

**Training, Validation and Test sets**

Glial cells were considered part of the positive class and non-glial cells (neuronal and anatomical) were considered as the negative cells. We construct the input expression matrix$X \in \mathbb{R}^{n_{cell} \times m_{gene}}$, containing the expression profiles of glia and non-glial cells belonging to glia or non-glial clusters, and a corresponding output label $y \in \mathbb{R}^{n_{cell} \times 1}$ denoting whether a cell is glia or non-glia using only hermaphrodite cells. This dataset was then randomly split into training (70%), validation (20%) and test (10%). The sampling was performed separately for each cluster to ensure representation of each cluster across all datasets.

**Classification between Glial and Non-Glial cells**

A logistic regression model was trained using **sklearn.linear_model.LogisticRegression (Pedregosa et al., 2012)** to predict class labels based on gene expression and is defined as follows|

$$P_{Glia}(x_{cell}) = \frac{1}{1 + e^{-(x_{cell}w + w_0)}}$$

Where:

- $x_{cell}$ is a vector of length <u>mgenes</u> where each entry is measured gene expression for a given cell
- $w \in \mathbb{R}^{m_{genes} \times 1}$ containing weights or parameter $w_i$ of the logistic regression model
- $w_0$ is a constant term
- $P_{Glia}(x_{cell})$ is the computed probability estimate of a given cell with expression profile$x_{cell}$ belonging to the positive class, glia

L1-lasso regularization penalty was used during training to ensure sparsity of selected features and generalization to test data. We define a set $C$ containing a range of regularization parameters (inverse of regularization strength parameter **C=0.001, 0.005, 0.01, 0.05, 0.1, 0.5, 1.0, 5.0**). We then optimize a set of models under these conditions as defined below and define the set of trained models corresponding to each regularization parameter $c_i \in C$ denoted by the following expression:

$$\min_{w} \quad J(y,\, P_{Glia}(x), c),\ \forall_c \in C$$

Where $J(y,\, P_{Glia}(x), c)$ is the log-loss cost function with L1 penalty defined as follows

$$\min_{w} \quad J(y,\, P_{Glia}(x), c)$$

$$= \sum_{i=1}^{n} \left[ y_i \log\left(P_{Glia}(x_i)\right) \; - \; (1 \; - \; y_i) \log\left(1 - P_{Glia}(x_i)\right) \right] + \frac{1}{c} \sum_{i=1}^{m} \left| |w_i| \right|_1$$

- $P_{Glia}(x_i)$ is the probability estimate for a given cell with gene expression vector $x_i$ the output of the logistic regression model
- $y_i$is the corresponding ground truth label for the given cell 
- $c$ the inverse of regularization parameter that is varied for each optimization, controlling how regularized the trained model will be
- $w_i$ is a parameter associated to a feature or gene input

L1 regularization effectively eliminates features that are not informative in predicting class labels and retains features that are informative for making predictions. Following training, performance was evaluated for each model by computing the classification accuracy (using **score** method) on the validation set and a model was selected based on highest mean accuracy.

**Feature Ranking & Selection**

We develop an iterative feature ranking procedure to identify the most informative features that can confidently identify all glial cell-types and thus represent a core set of pan-glial markers. We first rank individual genes by their ability to distinguish glial and non-glial cells by zeroing out the weights of the model associated with a gene except for a single weight (gene). We formalize this approach as follows

$w^{(g_i)}$: $w \, \exists_{w_i} = 0, \, \forall_{i \neq g_i}$ Where:

- $w$ is the set of weights used by the baseline model
- $w_i$ is the $i^{th}$ parameter associated with the $i^{th}$ gene that is set to zero in the model and is synonymous to gene being used for prediction
- $w^{(g_i)}$ is the new set of weights for the model associated with the $g^{th}$ gene of interest, where every weight is set to zero except for the $g^{th}$ index

To rank the individual genes, we use the set of modified weights $w^{(g_i)}$ to compute the probability estimate for each cell $x_i \in X$ and denote the results as $P_{Glia}^{(g_i)}(x_i) \in \mathbb{R}^{n_{cell} \times 1}$ as the probability estimates for each cell, $x_i$, when only the $g^{th}$ gene of interest is used to make predictions. Subsequently, we then compute the score as follows:

$$GeneScore(g_i) = \frac{1}{n_{Glia}} \sum_{i \in I}^{n_{glia}} P_{Glia}^{(g_i)}(x_i)$$

Where:

- $I$ is the set of all cells in the dataset that are Glia

- $P_{Glia}^{(g_i)}(x_i)$ is the computed probability estimate for the ith glial cell using the set of weights $w^{(g_i)}$
- $n_{Glia}$ Is the total number of glial cells in the dataset
- $g_i$ is the gene of interest for which a score is to be calculated, $g_i \in G$ where G$G$ is a set containing all the genes used for the model

This process is repeated for all genes used in the model and a gene $g_i \in G$ is selected that maximizes the computed score defined below:

$$\arg\max_{g \in G}[GeneScore(g)] = \arg \max_{g \in G}\left[\frac{1}{n_{Glia}}\sum_{i=1}^{n_{Glia}} P_{Glia}^{(g)}(x_i)\right]$$

We make note of this gene and establish a set $S$ containing the current set of selected genes. Using only this set of gene we compute the average probability estimate for each glial cluster in the data and identify the glial cluster, $k_i$, that the selected genes in set $S$ fails to confidently identify as glia. We formulate this process as follows:

$$\arg\min_{k \in K}[ClusterScore(g_i, k)] = \arg\min_{k \in K}\left[\frac{1}{n_{cell}}\sum_{i \in k}^{n_{cell}} P_{Glia}^{(g_i)}(x_i)\right]$$

- $K$ is the set of glia clusters in the dataset, which $k_i \in K$
- $k_i$ contains several cells $n_{cell}$ with each cell having expression profile $x_i$
- $g_i \in S$ and is the first gene selected gene

We then further select features in a iterative manner, in which we test a gene $g_i'$ in combination with the current set of genes $S$ such that the added gene, $g_i'$, should be able improve on glial cluster, $k_i$ that the previous set of gene$S$ had difficulty identifying as glia. To do this, we devise a way to zero every coefficient of the model except for a few weights as follows, extending our previously described method:

$$w^{(S_{g'})}: w \exists_{w_i} = 0, \forall_{i \notin S_{g'}}$$

Where:

- $w$ is the set of weights used by the baseline model
- $S_{g'}$ : $S \cup \{g_i'\}$ are the set of genes to be tested, where $g_i' \notin S$ and $g_i' \in G$ which is the gene of interest to be added to the currently selected set of genes $S$
- $w^{(S_{g'})}$ Is the new set of weights associated with the $g_i'^{th}$ gene of interest where all weight indices not specified in the set $S_{g'}$ are set to zero

To identify the next gene to be added we computed the average probability estimate for a previously identified cluster $k_i$, where the set of genes $S$ had difficulty identifying as glia, using the new weights $w^{(S_{g'})}$ associated with gene, $g_i'$, where we select for $g_i'$ that maximizes the mean probability estimates for cluster $k_i$ denoted as follows below:

$$\arg\max_{g' \in G}\left[GeneClusterScore(S_{g'}, k_i)\right] = \arg\max_{g' \in G}\left[\frac{1}{n_{cell}}\sum_{i \in k_i}^{n_{cell}} P_{Glia}^{(S_{g'})}(x_i)\right]$$

The selected gene is then added to set $S$ and a cluster, $k_i$, where the set of current genes performs poorly is therefore identified accordingly and is used as a constraint for the next gene selection.

$$\arg\min_{k \in K}\left[ClusterScore(S,\ k_i)\right] = \arg\min_{k \in K}\left[\frac{1}{n_{cell}}\sum_{i \in k_i}^{n_{cell}} P_{Glia}^{(S)}(x_i)\right]$$

In general, for the subsequent genes to be selected, the previous gene selection followed by identification of low performing clusters was repeated several times until a specified number of genes were selected. With each iteration of this analysis, the previously identified baseline model was modified such that all but a subset of specified features are ablated.

**Sheath & Socket glia**

Sheath and socket annotations were determined using hierarchical clustering. Mean of batch corrected principal components for each cluster were used to determine the pairwise distances between each pair of clusters. The pairwise distance matrix was hierarchical clustered using the **scipy.cluster.hierarchy** function (**linkage='average'** and **metric='euclidean'**). This resulted in two distinct clades which were annotated as sheath and socket based on annotation of glial cell-types. The distinction is consistent when the analysis was restricted to using only hermaphrodite or male clusters.

Figure S1

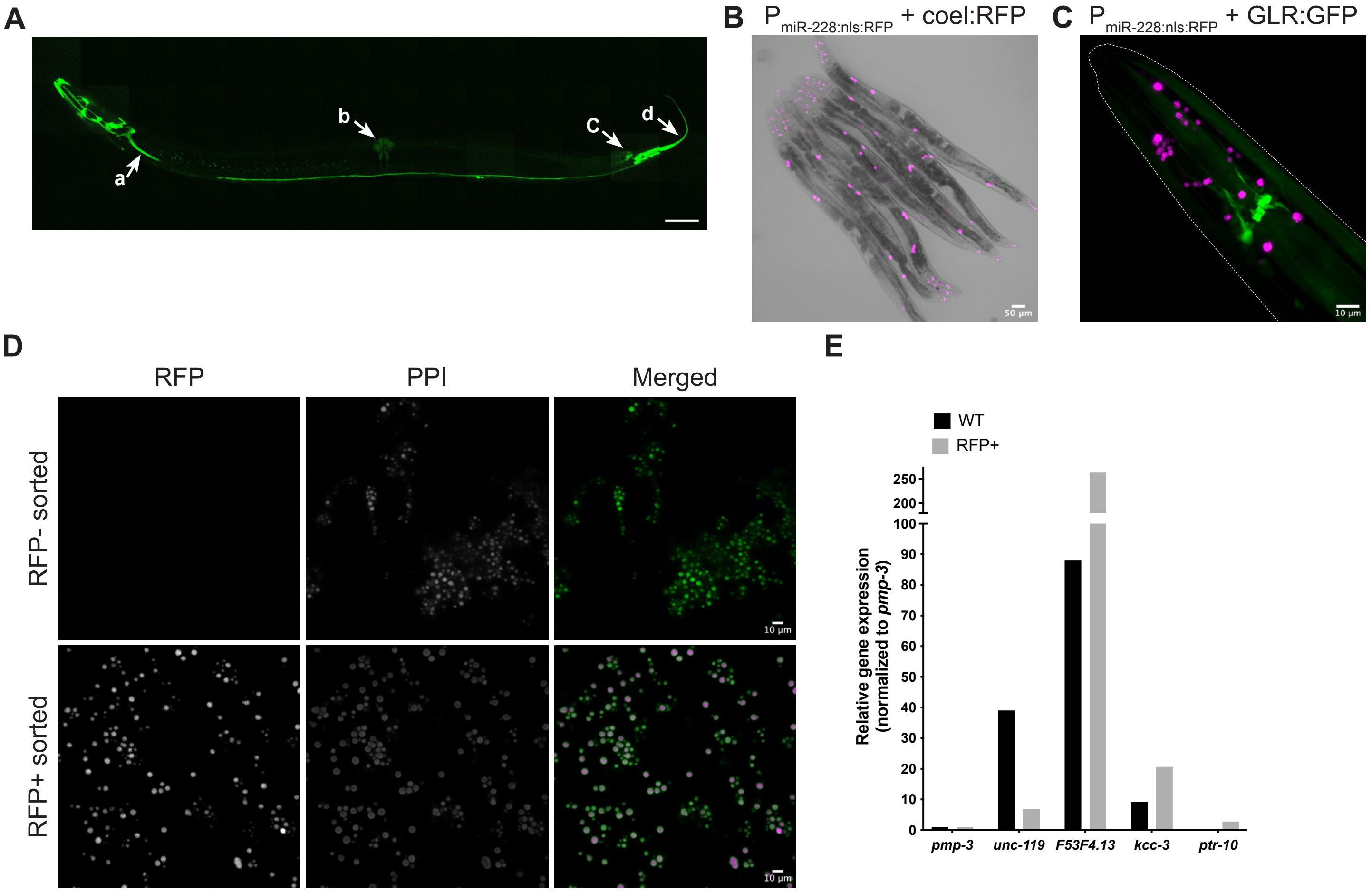

A
a
b
C
d
B
$P_{miR-228:nls:RFP}$ + coel:RFP
50 µm
C
$P_{miR-228:nls:RFP}$ + GLR:GFP
10 µm
D
RFP
PPI
Merged
RFP- sorted
RFP+ sorted
10 µm
10 µm
E
WT
RFP+
Relative gene expression (normalized to pmp-3)
250
200
100
90
80
70
60
50
40
30
20
10
0
pmp-3
unc-119
F53F4.13
kcc-3
ptr-10

Figure S2

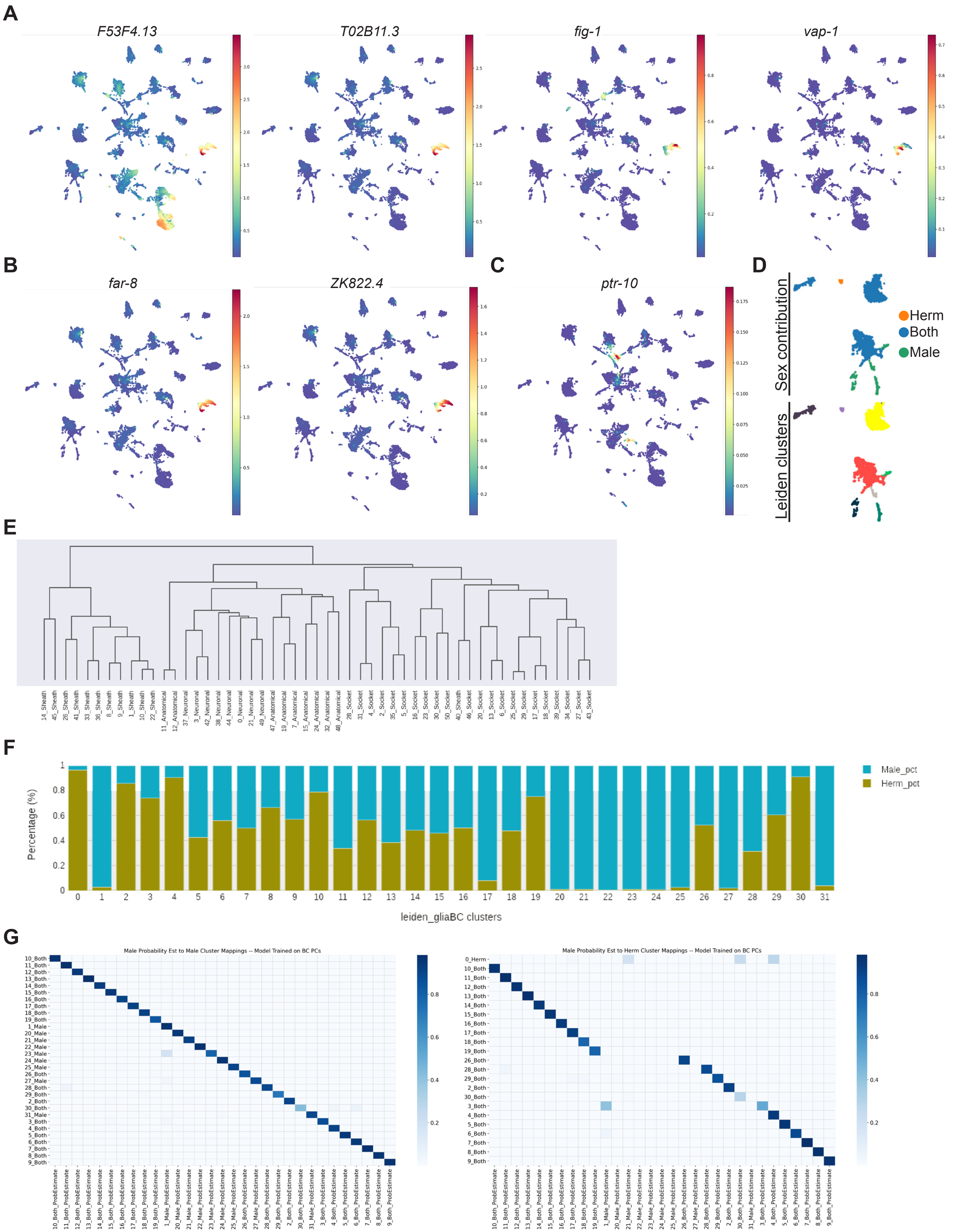
A
F53F4.13
T02B11.3
fig-1
vap-1
B
far-8
ZK822.4
C
ptr-10
D
Sex contribution
Leiden clusters
Herm
Both
Male
E
F
Percentage (%)
leiden_gliaBC clusters
Male_pct
Herm_pct
G
Male Probability Est to Male Cluster Mappings -- Model Trained on BC PCs
Male Probability Est to Herm Cluster Mappings -- Model Trained on BC PCs

Figure S2 continued

**H**

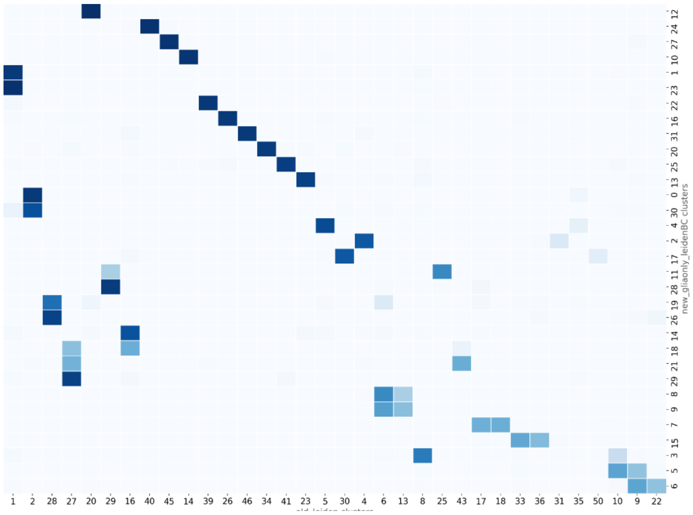


**I**

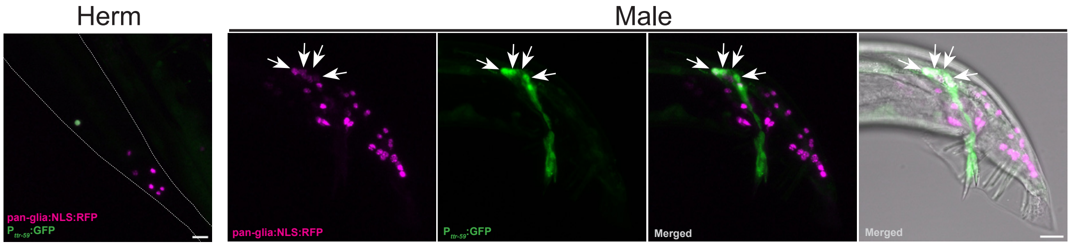


**J**

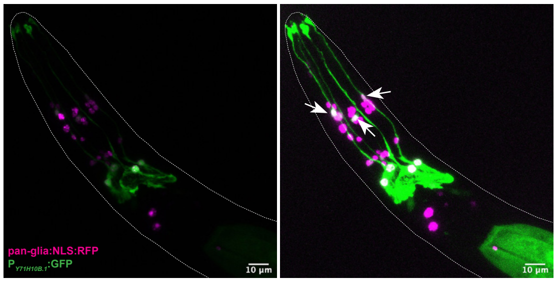

### Figure S3

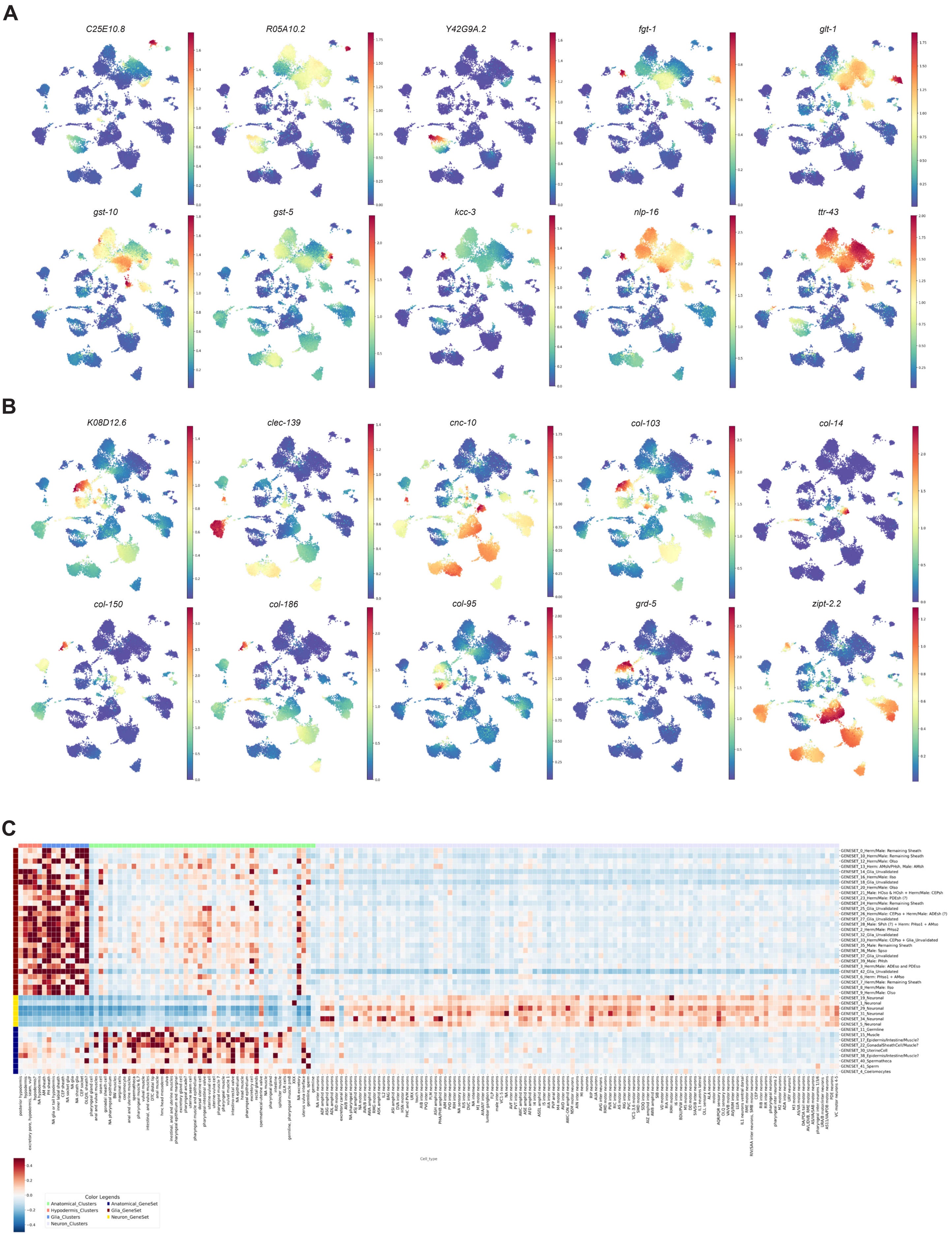
A
C25E10.8
R05A10.2
Y42G9A.2
fgt-1
glt-1
gst-10
gst-5
kcc-3
nlp-16
ttr-43
B
K08D12.6
clec-139
cnc-10
col-103
col-14
col-150
col-186
col-95
grd-5
zipt-2.2
C
GENESET_0_Herm/Male: Remaining Sheath
GENESET_10_Herm/Male: Remaining Sheath
GENESET_12_Herm/Male: OIso
GENESET_13_Herm: AMsh/PHsh, Male: AMsh
GENESET_14_Glia_Unvalidated
GENESET_16_Herm/Male: IIso
GENESET_18_Glia_Unvalidated
GENESET_20_Herm/Male: OIso
GENESET_21_Male: HOso & HOsh + Herm/Male: CEPsh
GENESET_23_Herm/Male: PDEsh (?)
GENESET_24_Herm/Male: Remaining Sheath
GENESET_25_Glia_Unvalidated
GENESET_26_Herm/Male: CEPso + Herm/Male: ADEsh (?)
GENESET_27_Glia_Unvalidated
GENESET_28_Male: SPsh (?) + Herm: PHso1 + AMso
GENESET_2_Herm/Male: PHso2
GENESET_32_Glia_Unvalidated
GENESET_33_Herm/Male: CEPso + Glia_Unvalidated
GENESET_35_Male: Remaining Sheath
GENESET_36_Male: Spso
GENESET_37_Glia_Unvalidated
GENESET_39_Male: PHsh
GENESET_3_Herm/Male: ADEso and PDEso
GENESET_42_Glia_Unvalidated
GENESET_6_Herm: PHso1 + AMso
GENESET_7_Herm/Male: Remaining Sheath
GENESET_8_Herm/Male: IIso
GENESET_9_Herm/Male: OIso
GENESET_19_Neuronal
GENESET_1_Neuronal
GENESET_29_Neuronal
GENESET_31_Neuronal
GENESET_34_Neuronal
GENESET_5_Neuronal
GENESET_11_Germline
GENESET_15_Muscle
GENESET_17_Epidermis/Intestine/Muscle?
GENESET_22_GonadalSheathCell/Muscle?
GENESET_30_UterineCell
GENESET_38_Epidermis/Intestine/Muscle?
GENESET_40_Spermatheca
GENESET_41_Sperm
GENESET_4_Coelomocytes
Cell_type
Color Legends
Anatomical_Clusters
Anatomical_GeneSet
Hypodermis_Clusters
Glia_GeneSet
Glia_Clusters
Neuron_GeneSet
Neuron_Clusters

Figure S3 continued

**D**

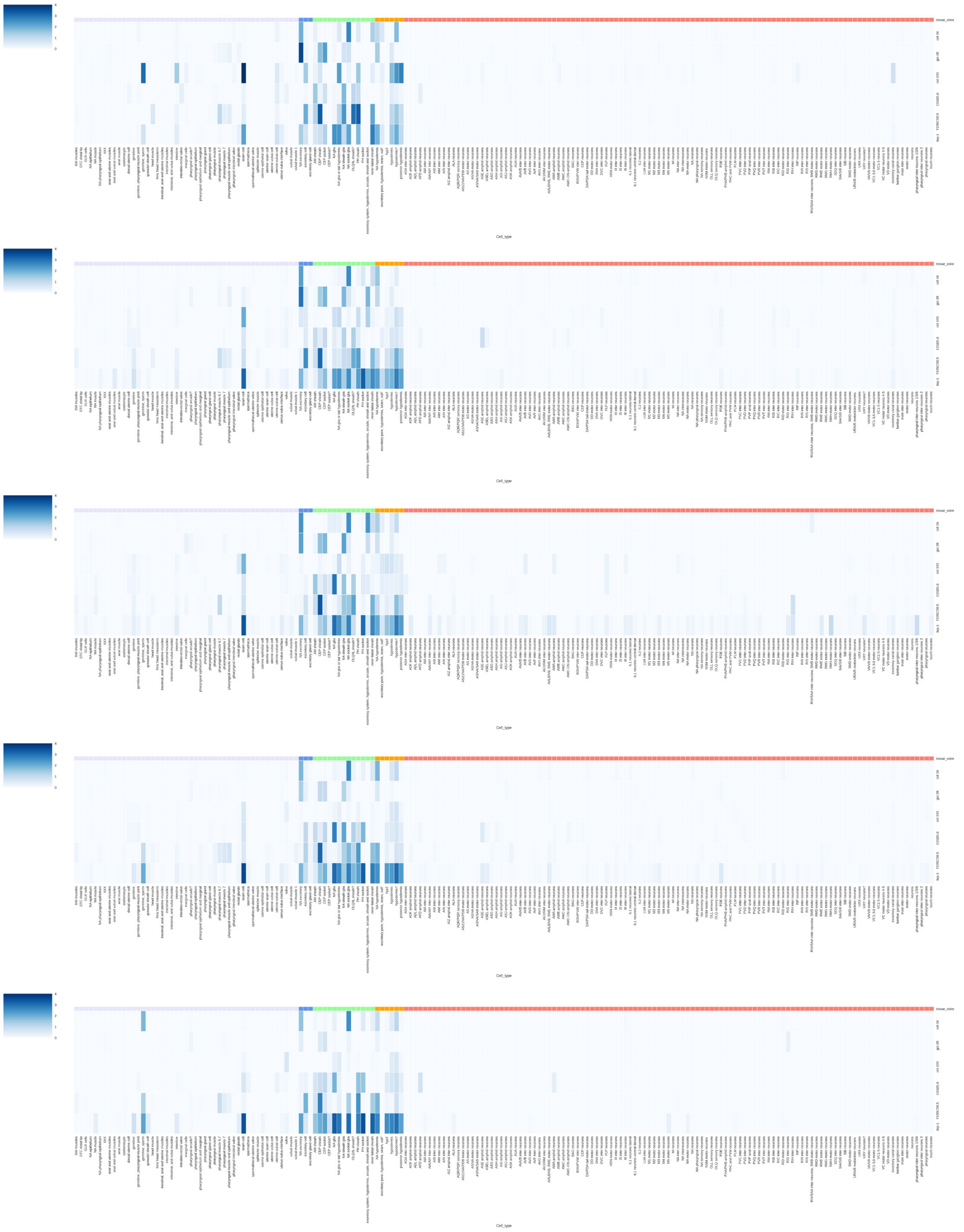

Figure S3 continued

**D continued**

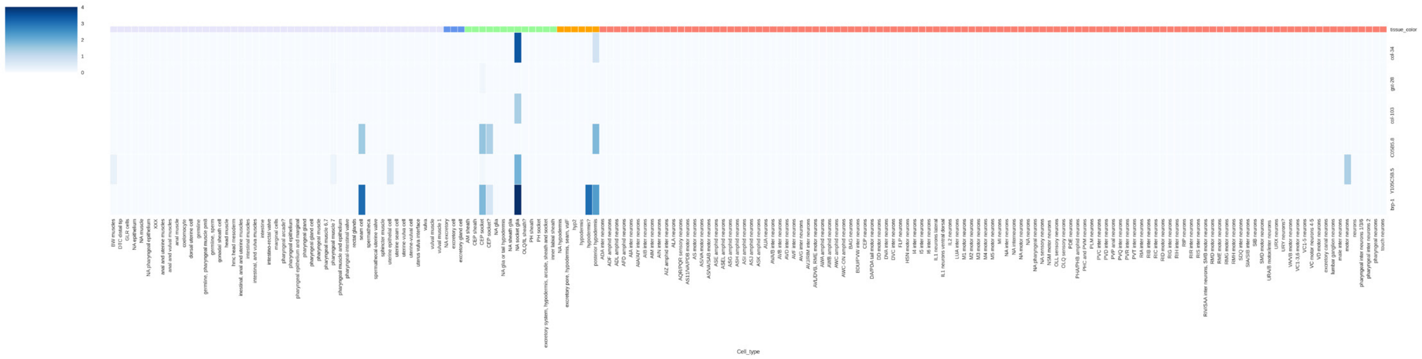

Figure S4

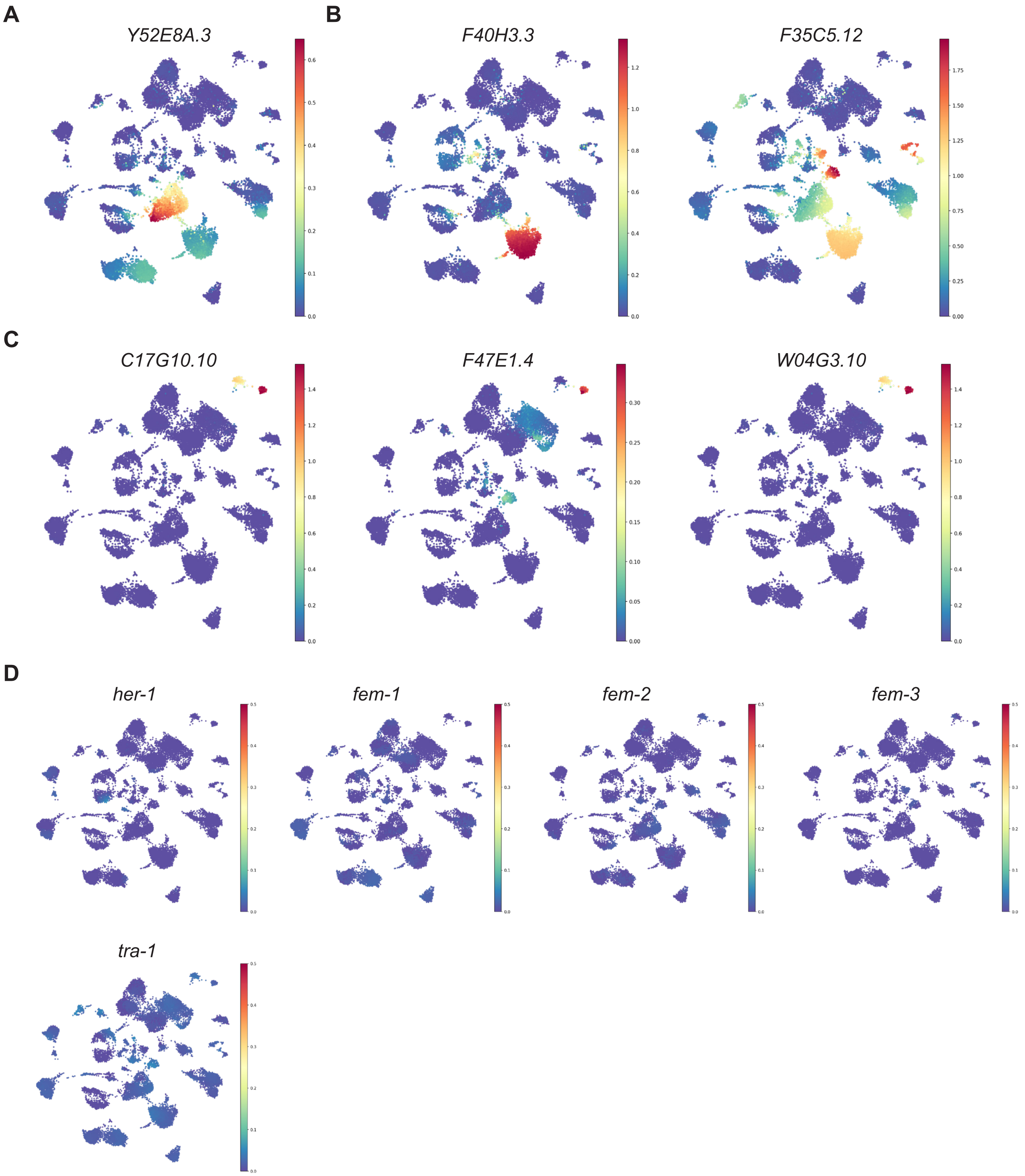
A
B
Y52E8A.3
F40H3.3
F35C5.12
C
C17G10.10
F47E1.4
W04G3.10
D
her-1
fem-1
fem-2
fem-3
tra-1

Figure S5

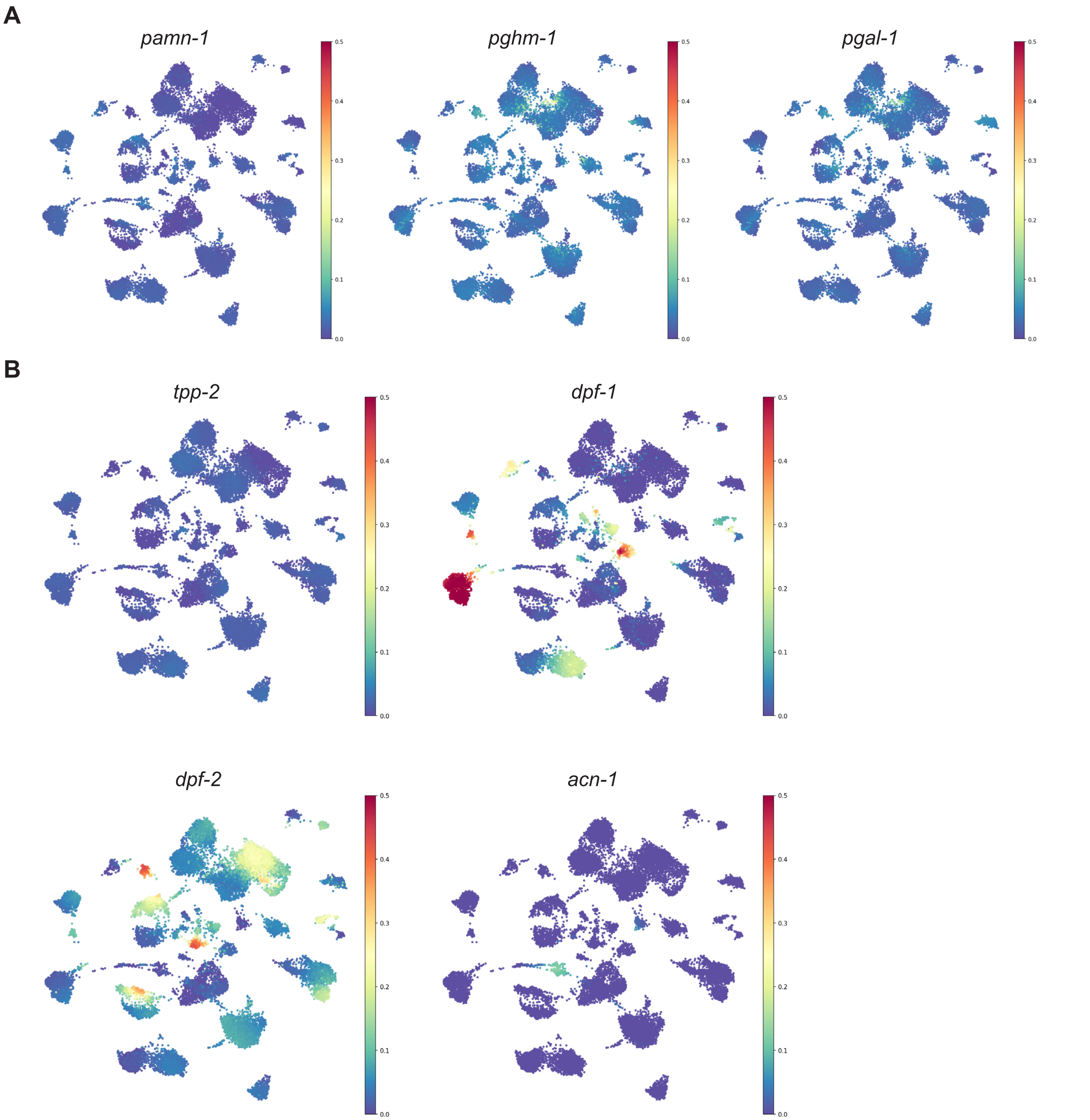

**SUPPLEMENTAL FIGURE LEGENDS:**

**Figure S1. Characterization of adult *C.elegans* glia using pan-glial transcriptional reporter *miR-228* and dissociation/nuclear sorting validations.** (**A**) Z-stack projection and stitched tiles depicting adult hermaphrodite expressing pan-glial cytoplasmic GFP. Arrows denote non-specific expression in (**a**) excretory canal, (**b**) vulva, (**c**) rectum, and (**d**) hypodermis. Scale bar = 50μM. (**B**) Z-stack projection of adult hermaphrodites expressing pan-glial nuclear RFP and co-injection marker coelomocyte RFP. No pan-glial nuclear RFP expression observed within the embryos of the germline. Scale bar = 50μM. (**C**) Z-stack projection of adult hermaphrodite head showing that GFP+ GLR glia does not expression pan-glial nuclear RFP. Scale bar = 10μM. (**E**) Single one-micron images of RFP- and RFP+ sorted nuclei after FACS stained with DNA die propidium iodide (PPI). Scale bars = 10μM. (**E**) Quantitative real-time PCR of selected neuronal (*unc-119*) and glial (*F53F4.13, kcc-3, ptr-10*) genes in RFP+ sorted nuclei and dissociated WT animals. Genotypes: Figure S1A: $P_{miR-228}$*:GFP*. Figures S1B-C: $P_{miR-228}$*:nls:RFP,* $P_{unc-122}$*:RFP*. Figures S1D*: him-5;* $P_{lgc-55}$*:GFP;* $P_{miR-228}$*:nls:RFP,* $P_{unc-122}$*:RFP*. Figure S1E, F: $P_{miR-228}$*:nls:RFP,* $P_{unc-122}$*:RFP* and *N2*.

**Figure S2. Validation of cluster identities via computational methods and transcriptional reporter validations.** (**A**) Expression of AMsh/PHsh genes (*F53F4.13, T02B11.3, fig-1, vap-1)* within the entire UMAP. (**B**) Expression of novel genes *far-8* and *ZK822.4* within the entire UMAP. (**C**) Expression of *ptr-10* within the entire UMAP. (**D**) The neuronal clusters and the sex-specificity of each cluster from the non-batch corrected UMAP. (**E**) Unbiased hierarchical clustering on the three subclusters of cells (glia split into sheath or socket, neurons, anatomical).

(**F**) Percentage of sex-specific cell contribution to each cluster. Cyan = male cells. Mustard = hermaphrodite cells. (**G**) Left: Machine learning model trained on the male dataset shows 1 to 1 maping to the manually assigned sex labels in (F). Right: With the exception of cluster 1, 3, and 30, machine learning model trained on the hermaphrodite dataset shows 1 to 1 mapping to the manually assigned sex labels in (F). (**H**) Pairwise comparison analyses between glia only batch-corrected (y-axis) and non-batch corrected datasets (x-axis) shows that some clusters from the non-batch corrected dataset merge in the batch corrected dataset. (**I**) Z-stack projections of the *ttr-59* transcriptional reporter. Hermaphrodite tail shown as merged. Male tail shown as individual and merged channels. Arrows point to the four *miR-228* RFP+ nuclei that co-localize with the GFP+ cells projecting into the spicule. (**J**) Z-stack projection of the *Y71H10B.1* transcriptional reporter in hermaphrodite head shows expression within the CEPsh glia. Image on right is the same image with increased brightness. Arrows depict CEPso glia expression. Genotypes: Figure S2I: *him-5;* $P_{ttr\text{-}59}$*:GFP,* $P_{unc\text{-}122}$*:GFP;* $P_{miR\text{-}228}$*:nls:RFP,* $P_{unc\text{-}122}$*:RFP*. Figure S2J: *him-5;* $P_{Y71H10B.1}$*:GFP,* $P_{unc\text{-}122}$*:GFP;* $P_{miR\text{-}228}$*:nls:RFP,* $P_{unc\text{-}122}$*:RFP*. All scale bars = 10μM.

**Figure S3. Unsupervised clustering identifies markers for populations of glia.** (**A**) Glial only batch corrected UMAP showing gene expression on top 10 sheath markers. (**B**) Glial only batch corrected UMAP showing gene expression on top 10 socket markers (**C**) Z-score analysis of the clusters in this dataset compared to those in Roux AE et al., 2022. (**D**) Pan-glial genes signatures scores across animal age in Roux AE et al., 2022. Expression for genes *col-34, gst-28, col-103, C0585.8, Y105C58.5,* and *brp-1* shown from top to bottom at each age.

**Figure S4. UMAP projections of sexually dimorphic genes and sex-determination genes.** (**A**) Glial only batch corrected UMAP showing *Y52E8A.3* gene expression. (**B**) Glial only batch corrected UMAP showing *F40H3.3* and *F35C5.12* gene expression. (**C**) Glial only batch corrected UMAP showing *C17G10.10, F47E1.4* and *W04G3.10* gene expression in male PHSh cluster (top right). (**D**) Glial only batch corrected UMAP showing lack of *her-1, fem-1/2/3, tra-1* gene expression in glial clusters.

**Figure S5. UMAP projections of genes related to neuropeptide processing.** (**A**) Glial only batch corrected UMAP showing expression levels of amidation enzymes (*pamn-1, pghm-1, pgal-1*). (**B**) Glial only batch corrected UMAP showing expression levels of multiple degradation enzymes (*tpp-2, dpf-1, dpf-2, acn-1*), half of which have high and variable expression.

**REAGENTS:**

Extrachromosomal array strains generated in this study (unless otherwise noted in the source).

| Strain | Source | Identifier |
|---|---|---|
| *him-5 V; nsIs708[PmiR-228:NLS-RFP + coel::RFP]* | This study | ASJ857 |
| *nsIs708 [PmiR-228:NLS-RFP + coel::RFP] ?* | Sean Wallace | OS11514 |
| *nsIs65[mig-24:GFP]; nsIs708 [miR-228:NLS:RFP + coel:RFP]* | This study | ASJ457 |
| *him-5(e1490?) V* | CGC | CB4088 |
| *nsIs198[Pmir-228:GFP]; nsIs708 [Pmir-228:NLS-RFP + coel:RFP]* | This study | ASJ445 |
| *him-5 V; nIs471 [lgc-55:GFP]; lin-15B & lin-15A(n765) X; nsIs708[Pmir-228:NLS-RFP + coel::RFP]* | This study | ASJ495 |
| *nsIs198[mir-228:GFP]* | This study | ASJ45 |
| *dnaEx235[Pfar-8:GFP + coel:GFP]; him-5 V; nsIs708[Pmir-228:NLS-RFP + coel::RFP]* | This study | ASJ765 |
| *dnaEx231[pZK822.4:GFP + coel:GFP]; him-5 V; nsIs708[Pmir-228:NLS-RFP + coel::RFP]* | This study | ASJ760 |
| *dnaEx246[PY67D8C.7.1:GFP + coel:GFP]; him-5 V; nsIs708[Pmir-228:NLS-RFP + coel::RFP]* | This study | ASJ767 |
| *dnaEx331[pY71H10b.1:GFP + coel:GFP]; him-5 V; nsIs708[Pmir-228:NLS-RFP + coel::RFP]* | This study | ASJ895 |
| *dnaEx331[pY71H10b.1:GFP + coel:GFP]; him-5 V; nsIs708[Pmir-228:NLS-RFP + coel::RFP]* | This study | ASJ776 |

| | | |
|---|---|---|
| *dnaEx451[Pzipt-2.2:GFP + coel GFP]; him-5 V; nsIs708[Pmir-228:NLS-RFP + coel::RFP]* | This study | ASJ1020 |
| *dnaEx395[pY52E8A.3:GFP + coel:GFP]; him-5 V; nsIs708[Pmir-228:NLS-RFP + coel::RFP]* | This study | ASJ890 |
| *dnaEx307[pF35C5.2:GFP + coel:GFP]; him-5 V; nsIs708[Pmir-228:NLS-RFP + coel::RFP]* | This study | ASJ781 |
| *dnaEx304[pF40H3.2:GFP + coel:GFP]; nsIs708 [Pmir-228:NLS-RFP + coel::RFP]; him-5 V* | This study | ASJ1022 |
| *dnaEx377[pCOL-177:GFP + coel:GFP]; him-5 V; nsIs708[Pmir-228:NLS-RFP + coel::RFP]* | This study | ASJ966 |
| *dnaEx305[pT27D12.1:GFP + coel:GFP]; nsIs708[Pmir-228:NLS-RFP + coel::RFP]* | This study | ASJ779 |
| *dnaEx247[Pttr-59:GFP + coel:GFP]; him-5 V; nsIs708[Pmir-228:NLS-RFP + coel::RFP]* | This study | ASJ778 |

Plasmids generated in this study.

| **PLASMID** | **Source** | **Identifier** |
|---|---|---|
| *PmiR-228:NLS-RFP* | This study | ASJ60/pMP15 |
| *Pfar-8:GFP* | This study | ASJ194/pMP44 |
| *pZK822.4:GFP* | This study | ASJ192/pMP42 |
| *pY67D8C.7.1* | This study | ASJ196/ pNT8 |

| | | |
|---|---|---|
| *pY71H10b.1* | This study | ASJ213/pNT21 |
| *Pzipt-2.2:GFP* | This study | ASJ348/pRSM18 |
| *pY52E8A.3:GFP* | This study | ASJ286/pNT53 |
| *pF35C5.2* | This study | ASJ217/pMP48 |
| *pF40H3.2* | This study | ASJ215/pMP46 |
| *pCOL-177:GFP* | This study | ASJ261/ pNT40 |
| *pT27D12.1* | This study | ASJ209/pNT17 |
| *Pttr-59:GFP* | This study | ASJ778/pNT7 |

**SUPPLEMENTAL REFERENCES:**


Kaletsky, R., Lakhina, V., Arey, R., Williams, A., Landis, J., Ashraf, J., Murphy, C.T., 2016. The C. elegans adult neuronal IIS/FOXO transcriptome reveals adult phenotype regulators. Nature 529, 92–96. https://doi.org/10.1038/nature16483

Korsunsky, I., Millard, N., Fan, J., Slowikowski, K., Zhang, F., Wei, K., Baglaenko, Y., Brenner, M., Loh, P., Raychaudhuri, S., 2019. Fast, sensitive and accurate integration of single-cell data with Harmony. Nat. Methods 16, 1289–1296. https://doi.org/10.1038/s41592-019-0619-0

McInnes, L., Healy, J., Saul, N., Großberger, L., 2018. UMAP: Uniform Manifold Approximation and Projection. J. Open Source Softw. 3, 861. https://doi.org/10.21105/joss.00861

Pedregosa, F., Varoquaux, G., Gramfort, A., Michel, V., Thirion, B., Grisel, O., Blondel, M., Müller, A., Nothman, J., Louppe, G., Prettenhofer, P., Weiss, R., Dubourg, V., Vanderplas, J., Passos, A., Cournapeau, D., Brucher, M., Perrot, M., Duchesnay, É., 2012. Scikit-learn: Machine Learning in Python. https://doi.org/10.48550/ARXIV.1201.0490

Taylor, S.R., Santpere, G., Weinreb, A., Barrett, A., Reilly, M.B., Xu, C., Varol, E., Oikonomou, P., Glenwinkel, L., McWhirter, R., Poff, A., Basavaraju, M., Rafi, I., Yemini, E., Cook, S.J., Abrams, A., Vidal, B., Cros, C., Tavazoie, S., Sestan, N., Hammarlund, M., Hobert, O., Miller, D.M., 2021. Molecular topography of an entire nervous system. Cell 184, 4329-4347.e23. https://doi.org/10.1016/j.cell.2021.06.023

Traag, V.A., Waltman, L., van Eck, N.J., 2019. From Louvain to Leiden: guaranteeing well-connected communities. Sci. Rep. 9, 5233. https://doi.org/10.1038/s41598-019-41695-z

van Dijk, D., Sharma, R., Nainys, J., Yim, K., Kathail, P., Carr, A.J., Burdziak, C., Moon, K.R., Chaffer, C.L., Pattabiraman, D., Bierie, B., Mazutis, L., Wolf, G., Krishnaswamy, S., Pe'er, D., 2018. Recovering Gene Interactions from Single-Cell Data Using Data Diffusion. Cell 174, 716-729.e27. https://doi.org/10.1016/j.cell.2018.05.061

Wolf, F.A., Angerer, P., Theis, F.J., 2018. SCANPY: large-scale single-cell gene expression data analysis. Genome Biol. 19, 15. https://doi.org/10.1186/s13059-017-1382-0
